# Suspension-Based Human Esophagoids Recapitulate WNT2B-Dependent Regulation of Esophageal Basal Progenitors

**DOI:** 10.64898/2026.06.23.733451

**Authors:** Noah Etzioni, Tristan Frum, Kelli Johnson, Andrea P. Álvarez-Maldonado, Helena M. Yllescas-Lopez, David E. Bayer, Zhiwei Xiao, Madeline K. Eiken, Claudia Loebel, Joshua H. Wu, Yu-Hwai Tsai, Angeline Wu, Charles J. Zhang, Michael K. Dame, Bhargav Gunuguntla, Ashley J Cuttitta, Hana Ho, Dominic J Tigani, Jonathan Sexton, Venkata S. Dasuri, Nicholas Makogonov, Amy E. O’Connell, Jason R. Spence, Daysha Ferrer Torres

## Abstract

**Background & Aims:** The human esophagus undergoes a tightly regulated developmental program, transitioning from a simple columnar epithelium in early development to a mature stratified squamous tissue essential for adult barrier function. Here, we constructed a developmental cell atlas spanning early development to adulthood and leveraged it to generate physiologically relevant *in vitro* models.

**Methods:** We utilized single-cell RNA sequencing and spatial multiplex proteomics of human esophageal tissue from early development through adulthood. We established a feeder-supported 2D culture system and a Matrigel-free, suspension-based 3D esophagoid model in a 96-well format. To interrogate WNT2B function, we analyzed patient tissue harboring WNT2B loss-of-function mutations and performed WNT inhibition in esophagoids.

**Results:** Sequencing profiling identified stage-specific epithelial populations: multiciliated and GPC3⁺ basal cells were unique to early development; KRT14⁺ basal and CRNN⁺ luminal cells were adult-specific; and COL17A1⁺, LY6D⁺, and KRT4⁺ populations were shared across stages. Spatially organized WNT2B⁺, KIT⁺, and VWC2⁺ mesenchymal subtypes were identified. The 2D system preserved both epithelial and mesenchymal compartments with transcriptional fidelity. Esophagoids exhibited basal-to-luminal stratification, mesenchymal compartmentalization, and required stromal interactions for formation. WNT2B repressed self-renewal of TP63⁺ basal progenitors and inhibited proliferation, confirmed by pharmacologic inhibition of WNT in the *in vitro* esophagoids.

**Conclusions:** We present a stage-resolved atlas of human esophageal development and a scalable esophagoid platform recapitulating esophageal architecture. WNT2B regulates progenitor dynamics by restraining basal cell self-renewal. Esophagoids provide a physiologically relevant system for modeling esophageal development and disease.

**Visual Abstract:** 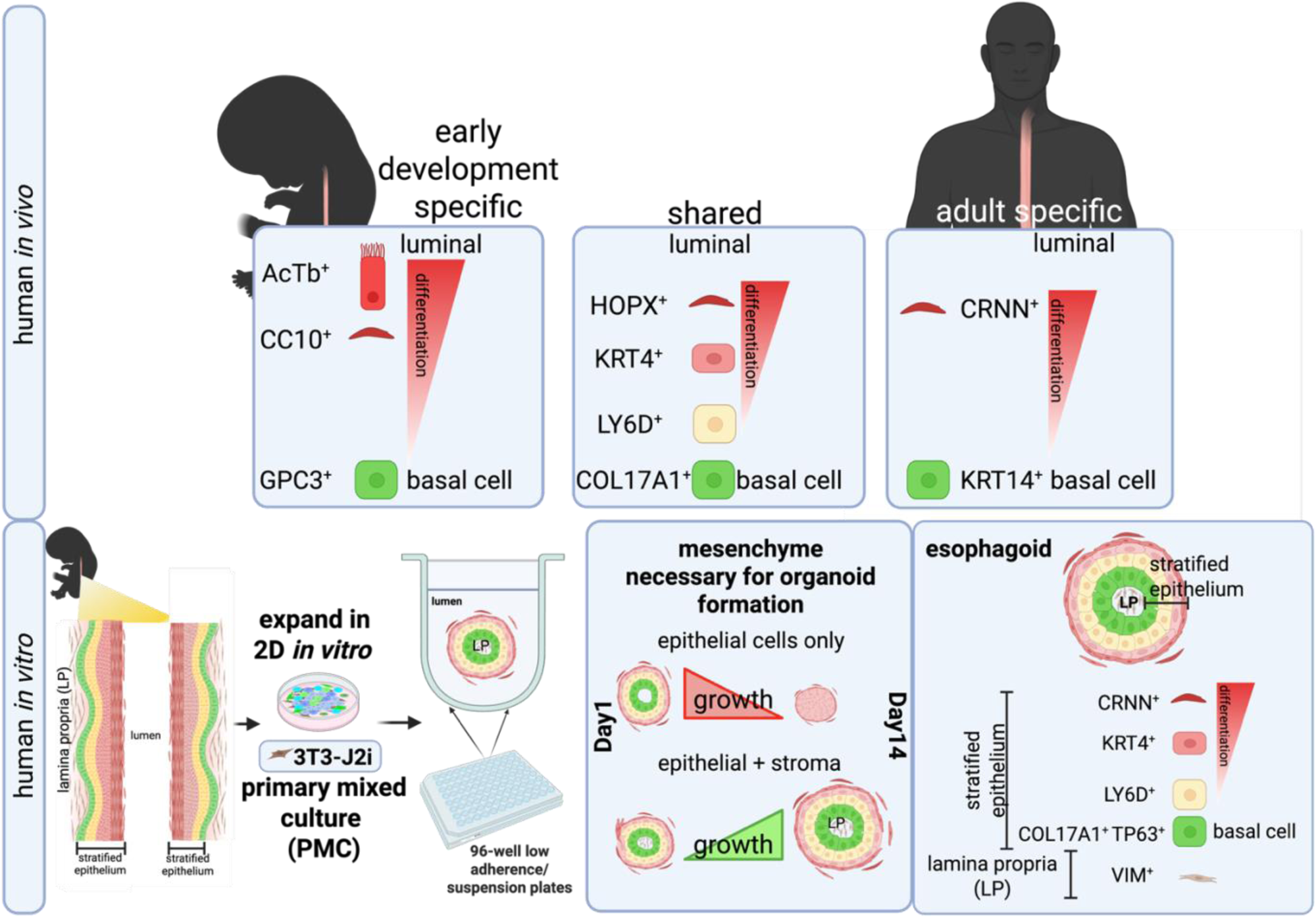

*Key Findings and Implications:*

- Developmental Atlas: The study presents a comprehensive transcriptional and structural atlas of the human esophageal epithelium, identifying conserved and stage-specific epithelial populations from early development to adulthood. Notably, stage-specific gene expression of multiciliated and *GPC3⁺* basal cells were unique to early development, while KRT14⁺ basal and CRNN⁺ luminal cells were adult specific, with COL17A1^+^ (basal), LY6D^+^ (epibasal), and KRT4^+^ (middle), shared at all stages.
- Mesenchymal Diversity: Spatial and transcriptional profiling revealed distinct mesenchymal subtypes, including *WNT2B*⁺, *KIT*⁺, and *VWC2*⁺ populations, which are spatially organized and contribute to epithelial-mesenchymal signaling. These findings reinforce the role of stromal-epithelial interactions in esophageal development.
- 2D Esophagus Cell Culture System: A feeder-supported 2D cell culture system was developed that retains both epithelial and mesenchymal populations, preserving transcriptional fidelity and enabling long-term expansion for mechanistic studies.
- 3D Esophagoid Model: A suspension-based 3D organoid system was optimized using a 96-well format, enabling high-throughput generation of esophagoids with robust epithelial stratification and mesenchymal compartmentalization. These organoids recapitulate key features of the human esophagus, including basal-to-luminal organization, and require stromal interactions for formation.
- Functional Role of *WNT2B* in esophagus development: Both *in vivo* and *in vitro* analyses demonstrated that WNT2B regulates epithelial progenitor dynamics and tissue architecture by repressing self-renewal of basally localized TP63+ cells and inhibiting proliferation. Loss-of-function models and WNT pathway modulation confirmed its role in epithelial-mesenchymal crosstalk and organoid integrity.

## Introduction

The esophagus is a dynamic organ that undergoes complex developmental processes to form a stratified squamous epithelium, which is capable of withstanding mechanical stress and protecting underlying tissues from environmental insults (Rosekrans, Baan et al. 2015, Zhang, Jiang et al. 2017, EŞRefoglu, Taslidere et al. 2018, Morrisey and Rustgi 2018). Development of the esophagus is tightly regulated by interactions between the epithelium and mesenchyme, which provide critical cues for cell proliferation, differentiation, and tissue architecture (Goss, Tian et al. 2009, Rodriguez, Da Silva et al. 2010, Zhang, Yang et al. 2018). Classic histological studies indicate that the human esophagus originates from the dorsal anterior foregut and becomes separated from the ventral trachea between approximately 4 and 6 weeks of gestation (Zhang, Bailey et al. 2021). At 8 weeks of gestation, the esophagus is lined by a pseudostratified columnar epithelium (DeNardi 1991). By approximately five months of development, the stratified squamous epithelium first appears in the middle third of the esophagus and progressively extends toward both the rostral and caudal ends, replacing the ciliated epithelium (Johns 1952). The esophageal epithelium transitions from a simple, proliferative state to a fully stratified tissue composed of basal progenitor cells, proliferative intermediates, and differentiated luminal cells (Geboes 1978, Huff and Carreon 2019, Edwards, Shacham-Silverberg et al. 2021). This process is tightly regulated by epithelial-mesenchymal interactions, which provide critical cues for cell proliferation, differentiation, and tissue architecture (Yang, McCullough et al. 2025). Most of our understanding of these interactions, have come from studying the mouse esophagus (Zhang, Jiang et al. 2017, EŞRefoglu, Taslidere et al. 2018, Zhang, Bailey et al. 2021), nonetheless, this organ holds striking differences between species (Grommisch, Eenjes et al. 2025). For example, the adult mouse esophageal epithelium is a three-to-four cell layer thick squamous keratinized epithelium (Rosekrans, Baan et al. 2015), while the human esophageal epithelium is up to 40 cell layers thick and displays prominent stromal invaginations, called papillae (Huff and Carreon 2019), in addition to submucosal glands (Long and Orlando, 1999), both of which are absent in mouse.

Advances in single-cell transcriptomics and spatial profiling have begun to reveal the cellular heterogeneity and developmental trajectories within the gastrointestinal tract (Madissoon, Wilbrey-Clark et al. 2019, He, Wang et al. 2020, Busslinger, Weusten et al. 2021, Chen, Zhao et al. 2021, Yu, Kilik et al. 2021, Ferrer-Torres, Wu et al. 2022, Rochman, Wen et al. 2022, Ding, Garber et al. 2024, Yang, McCullough et al. 2025). More recent efforts have focused on characterizing both the developing (Yu, Kilik et al. 2021, Yang, McCullough et al. 2025) and adult human esophagus (Busslinger, Weusten et al. 2021, Ferrer-Torres, Wu et al. 2022, Rochman, Wen et al. 2022). However, at this point in time, a comparison between the developing human esophagus epithelium and the mature adult esophagus epithelium has not been carried out. Therefore, to investigate human esophageal development, we utilized publicly available single-cell RNA sequencing datasets spanning both early developmental stages and adult esophageal epithelium (Yu, Kilik et al. 2021, Ferrer-Torres, Wu et al. 2022). This analysis identified stage-specific markers and epithelial subtypes, providing new insights into the dynamic cellular organization and lineage transitions between developing and adult human esophagus previously unknown.

Single-cell and spatial transcriptomics have revealed the intricate cellular architecture of the developing human stromal esophagus populations. Yang et al. (2025) presented a spatiotemporal multi-omics census that identified fibroblast progenitors and stratified mesenchymal subtypes organized in concentric layers around the epithelium. Building on this framework, our study further defines and characterizes the mesenchymal populations during early human esophageal development, identifying WNT2B⁺, KIT⁺, and VWC2⁺ mesenchyme populations with distinct spatial localization.

Finally, advances in organoid technology have been made to generate esophageal organoids from human pluripotent stem cells, with a major focus on the epithelial components of these organoids (Trisno, Philo et al. 2018, Zhang, Yang et al. 2018, Yang, McCullough et al. 2025). Additionally, multiple methods to generate epithelial organoids from primary human tissues have also led to complex epithelial organoid models (Giroux, Lento et al. 2017, Kasagi, Chandramouleeswaran et al. 2018, Ferrer-Torres, Wu et al. 2022, Milne, Mustafa et al. 2024). Here, we build on our prior methods developed to culture adult esophagus from biopsies (Ferrer-Torres, Wu et al. 2022) to generate long-term 2D cocultures from early developmental human esophagus, with the ability to retain both epithelial and mesenchymal populations *in vitro* over long term culture. We further develop a high throughput 3D suspension-based esophagoid platform that also maintains the diversity of epithelial and mesenchymal populations. Using this *in vitro* esophagoid model system, we demonstrate that WNT signaling plays a critical role in epithelial progenitor regulation and tissue architecture, and we couple these functional experiments with characterization of *WNT2B* mutant patient esophagus and esophagus tissue, showing that esophagoids can be used to model human relevant disease phenotypes.

In this study, we characterize the human esophagus establishing new mesenchymal sub-types and establish a scalable, physiologically relevant esophagoid model that features a mesenchymal center, epithelial basal stem cells, and a stratified epithelium, enabling functional interrogation of epithelial-mesenchymal crosstalk in esophageal development and disease.

## Results

### Transcriptional and Structural Dynamics of Esophageal Epithelium: From Early Development to Adult Differentiation

Our first goal was to detail a complete atlas of the transcriptional, structural, and developmental dynamics of the esophageal epithelium during early development and in the adult stage. We utilized single cell RNA sequencing publicly available data to compare early development (Yu, Kilik et al. 2021) vs adult (Ferrer-Torres, Wu et al. 2022) (Supplemental Figure 1A-D). Utilizing this data, we were able to identify and map epithelial, mesenchymal, and immune components of the esophagus (Supplemental Figure 1B), and highlight the epithelial and mesenchymal populations, with *CDH1* (epithelial) and *VIM* (mesenchyme) (Supplemental Figure 1A, D). Our first goal was to assess and compare the epithelial cell types between human esophagus in early development and adult (Figure 1, Supplemental Figure 1E-F). UMAP embeddings of the adult and early development esophageal epithelium revealed distinct epithelial populations (Figure 1A-B). The adult epithelium is stratified into five different populations including basal, proliferative, epibasal, middle, and luminal populations (Figure 1A), consistent with previous observations from our group (Ferrer-Torres, Wu et al. 2022) and others (Busslinger, Weusten et al. 2021, Yu, Kilik et al. 2021, Rochman, Wen et al. 2022). In contrast, early developmental epithelium includes additional transitional states, including two proliferative clusters, epibasal, late, and multiciliated epithelial populations (Figure 1B). To spatially map and confirm the epithelial identity of the early human esophagus cell populations identified in the sequencing data, we performed immunofluorescence staining for key lineage markers (Figure 1C). We observed robust expression of TP63, a basal/epibasal marker, and LY6D, an epibasal-specific marker, alongside KRT4, which marks middle-differentiation epithelial cells. These markers were previously characterized in the adult esophagus (Ferrer-Torres, Wu et al. 2022), indicating conserved expression patterns across developmental stages. Notably, the early developmental luminal epithelium lacked expression of CRNN, a marker of terminally differentiated luminal cells found in the adult esophagus (Figure 1A–C), suggesting a less mature epithelial state. Additionally, we detected expression of acetylated tubulin (AcTB), a marker of ciliated cells, a specialized epithelial subtype known to be present during early esophageal development (Linda M. Ernst and Eduardo D. Ruchelli , Menard 1995, Yaron Daniely 2004) (Figure 1C), and not found in adult samples (Supplemental Figure 1G). Label transfer analysis highlighted the developmental trajectory, showing alignment between early development and adult clusters, with basal and proliferative early development clusters corresponding to basal and early/epibasal adult populations (Figure 1D). Analysis of key genes across epithelial clusters in both adult and early developmental revealed shared and uniquely expressed genes (Figure 1E). These plots identified development-specific markers such as GPC3 and CC10, as well as shared markers like COL17A1, KRT4, LY6D, and HOPX, which persist into adulthood (Figure 1D, E, F; Supplemental Figure 1B). Notably, GPC3 was identified as an early development basal-specific marker, whereas KRT14, a well-established basal cell marker (Rosekrans, Baan et al. 2015), was exclusively expressed in adult basal cells (Figure 1F, G). Multiciliated markers were restricted to the early development stage (Figure 1A, B, D, E; Supplemental Figure 1G), while the cornified cell marker CRNN was solely expressed in adult luminal cells (Figure 1A, B, C). Histological analyses from early development stages to adulthood revealed progressive epithelial stratification, transitioning from a simple structure in early development to a fully differentiated, multilayered epithelium in adults (Supplemental Figure 1E, F).

**Figure 1.**
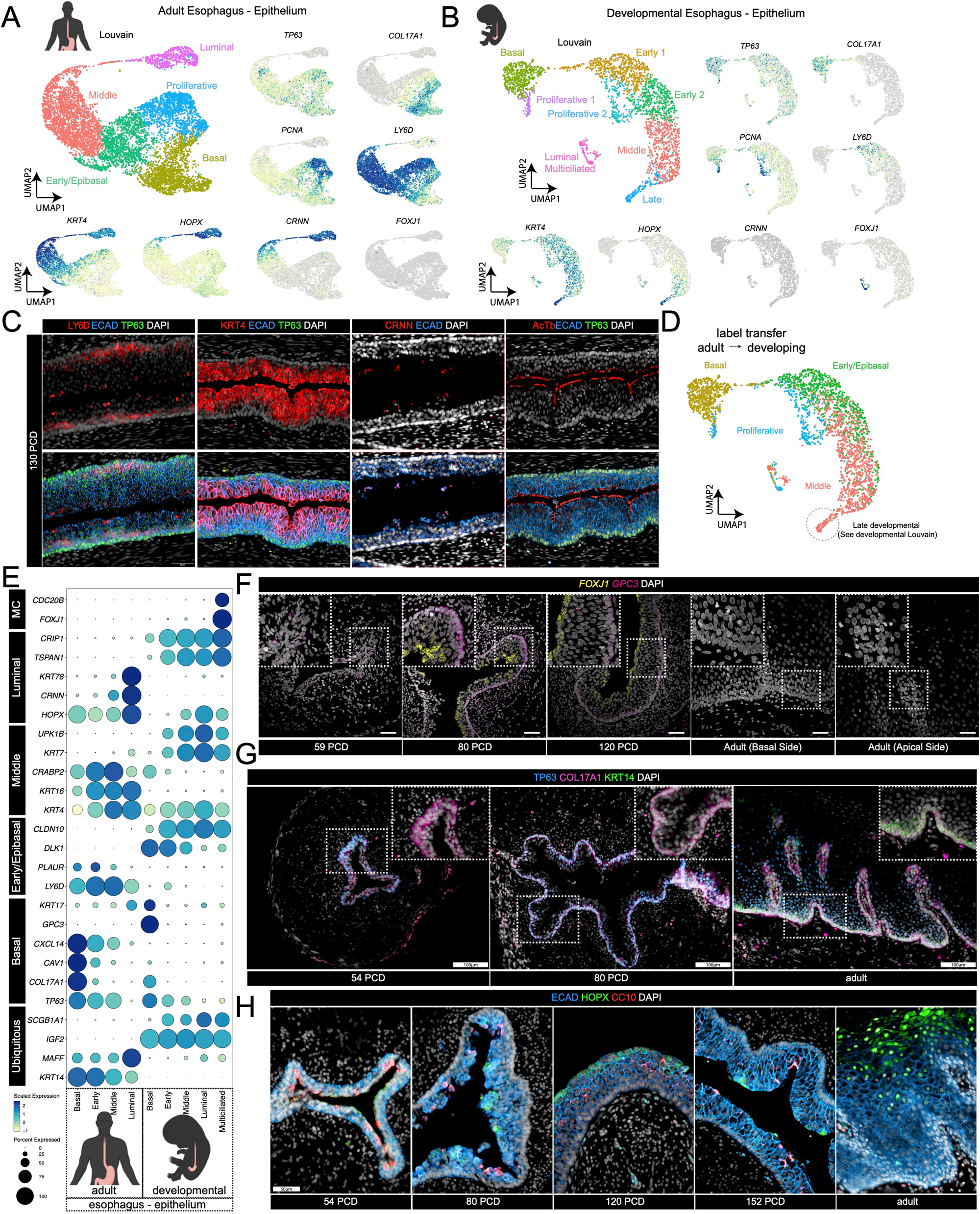
Comparative Analysis of Epithelial Cell Types in Adult and Early Development Human Esophagus Identifies Shared and Zone-Specific Markers. (**A**) Louvain clustering and UMAP visualization of epithelial cell populations in the adult human esophagus, highlighting distinct basal and luminal cell zones. (**B**) Louvain clustering and UMAP visualization of epithelial cell populations in the early development human esophagus, demonstrating comparable and unique cellular subtypes. (**C**) Immunofluorescence staining of early development (130 post conception days (PCD) esophageal tissue showing expression of LY6D, TP63, KRT4, CRNN, AcTB, nuclear marker DAPI and epithelial marker ECAD, confirming the presence of basal, epibasal, luminal epithelial cells. (**D**) Label transfer analysis mapping adult epithelial cell populations onto early development epithelial populations, revealing developmental relationships and conserved markers. (**E**) Dot plot of key gene expression patterns across epithelial clusters in both adult and early development esophagus, illustrating shared and cell zone-specific markers. **(F–H)** Immunofluorescence and fluorescence *in situ* hybridization analyses confirm the spatial distribution and expression of zone-specific markers in the basal and luminal epithelial layers of the adult and early development human esophagus.

Together, these data highlight both conserved architecture and stage-specific differences in the esophageal epithelium. We validated development-specific populations such as the basal cell-specific marker *GPC3*, multiciliated cells, and CC10 positive luminal cells from adult-specific populations, which expressed KRT14 restricted to the basal layer and CRNN positive cornified luminal cells. Notably, both developmental and adult stages shared expression of COL17A1 in basal stem cells, LY6D in epibasal cells, and KRT4 in mid to late differentiated epithelial cells (Supplemental Figure 1I). These findings collectively illustrate the coordinated cellular, transcriptional, and structural evolution of the esophageal epithelium, defining the cellular and molecular transitions accompanying esophageal epithelial maturation.

### Transcriptional Diversity and Spatial Organization of Mesenchymal Populations in Esophageal Development

We next wanted to characterize the mesenchymal populations present at different in the esophagus (Supplemental Figure 1A-B, Figure 2). Unfortunately, since the adult tissue was taken from biopsies, there was a very small mesenchymal population in this sample cohort (Supplemental Figure1 C-D), and comparisons between adult and development was not carried out (Figure 2). To explore the mesenchymal cell types, present in the developing esophagus and map their spatial organization, we first interrogated UMAP embeddings of mesenchymal cells, identifying distinct subtypes including: proliferative mesenchyme, *WNT2B*+ mesenchyme, *APOE*+ mesenchyme, *KCNN3*+ mesenchyme, *NRXN1*+mesenchyme, *KIT*+ mesenchyme, smooth muscle precursors, and mature smooth muscle (n=8 clusters) (Figure 2A). These clusters highlight the heterogeneity within the mesenchymal compartment. Figure 2B presents a dot plot of key genes expressed across these mesenchymal subtypes. Notable markers include *PDGFRA*, highly expressed in *WNT2B*+mesenchyme, *GDF10* in multiple populations, and *KIT*, which specifically identifies the *KIT*+ mesenchyme cluster (Figure 2B). From each subcluster subtype we focus on highly expressed genes, such as feature plots of genes *PDGFRA, GDF10, HGF, KIT, WNT2B, VWC2,* and *TOP2A*, which show unique spatial distributions within the UMAP. These genes define the identities of broad as well as specific mesenchymal clusters, with *WNT2B* and *HGF* specifically marking mesenchyme populations that are critical for epithelial-mesenchymal crosstalk.

**Figure 2.**
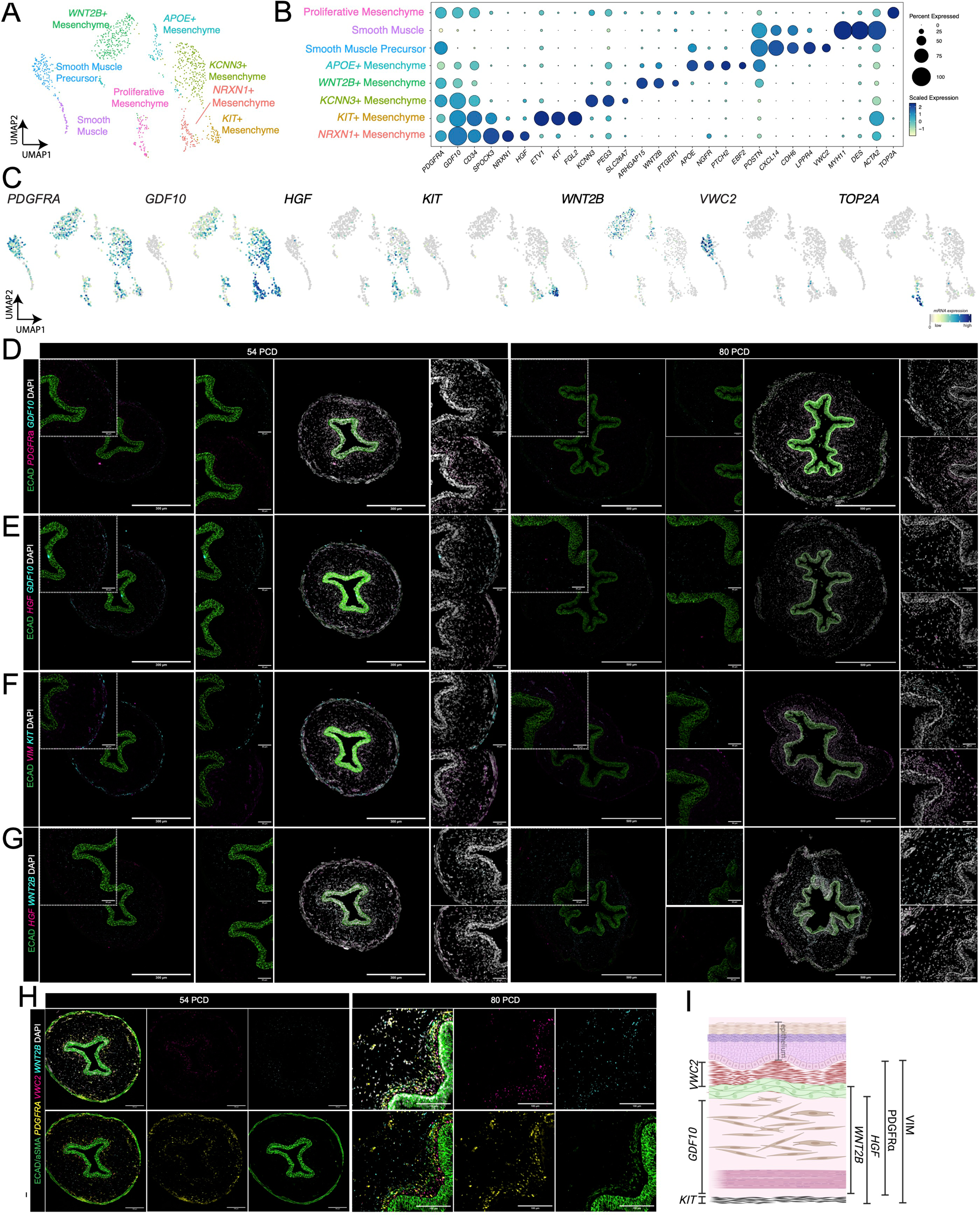
Comprehensive Characterization of Mesenchymal Cell Populations in the Human Esophagus during Early Development. **(A)** Louvain clustering and UMAP analysis of mesenchymal cell populations in the early development human esophagus, delineating distinct cellular subtypes. **(B)** Dot plot displaying the top differentially expressed genes for each identified mesenchymal cell population, highlighting key molecular markers. **(C)** UMAP projections visualizing the expression patterns of representative genes across mesenchymal cell clusters. **(D–H)** Immunofluorescence combined with fluorescence in situ hybridization (FISH) validates the spatial localization and expression of key genes within mesenchymal cell populations in the early development esophagus. **(I)** Summary diagram illustrating the gene expression profiles and spatial organization of mesenchymal cell subtypes in the early development human esophagus.

We proceeded to characterize and validate the spatial distribution and expression of top cluster defining genes, utilizing fluorescence in situ hybridization (FISH) coupled with immunofluorescence in sections at 54 and 80 days of development (Figure 2D-H). *PDGFRA, WNT2B, HGF*, and *KIT* are shown to colocalize independent of the epithelial marker ECAD, indicating their physical association close to, but not within, the developing epithelium. We validated by FISH that *PDGFRA* as well as *VIM* are expressed in all the mesenchymal populations (Figure 2D-F), as observed in the single cell data (Figure 2C). We observed that *GDF10* was found to be co-expressed in the *PDGFRA* population as well as within a subset of *HGF* cells (PDGFRA^+^/GDF10^+^/HGF^+^), nonetheless, there is a specific subset of *GDF10* that do not express *HGF (*PDGFRA^+^/GDF10^+^/HGF^-^) (Figure 2C-E) while *HGF* seems to be expressed in a more defined mesenchymal population by 80 days of development (Figure 2E-G). We also identified mesenchymal populations marked by *KIT*, *WNT2B,* and *VWC2* (Figure 2F, G, H). *VWC2* was specifically expressed in the mesenchymal population closest to the epithelium (Figure 2H), while *KIT* was expressed in the outer most layer of the mesenchyme (Figure 2F). Finally, co-staining of *PDGFRA* and *WNT2B* highlights overlapping yet distinct mesenchyme (Figure 2H), further demonstrating the spatial heterogeneity of these cells during development. Figure 2I provides a schematic summarizing the spatial distribution of mesenchymal populations within the developing human esophagus.

Together, these data reinforce the concept of spatially organized mesenchymal during esophageal development and highlight conserved stromal-epithelial interactions across studies (Yang, McCullough et al. 2025). These findings highlight the essential contributions of mesenchymal heterogeneity to the development of the esophageal epithelium and smooth muscle, providing a framework for understanding epithelial-mesenchymal interactions in both normal development and disease contexts.

### 2D In Vitro Expansion of Developmental Esophageal Cells Using a 3T3-J2 Feeder Cell System

To model esophageal development *in vitro*, human early-stage esophageal tissues were mechanically dissociated and expanded as a 2D monolayer on a layer of 3T3-J2 cells (n=3 at development stages 101, 122, and 130 post conception days) (Figure 3A). Representative brightfield microscopy revealed the successful establishment and propagation of epithelial and mesenchymal colonies, which progressively increased in size and density over successive passages, as shown at passage 1 (P1) and passage 3 (P3) (Figure 3B). Immunofluorescence staining for human nuclei (Hu-Nu) confirmed the human origin of the esophageal cells (Figure 3C). The 2D cultures consisted of both epithelial and mesenchymal cells, as demonstrated by immunostaining for ECAD (*CDH1*; epithelial marker) and VIM (mesenchymal marker), which revealed distinct spatial segregation of the two cell types (Figure 3D).

**Figure 3.**
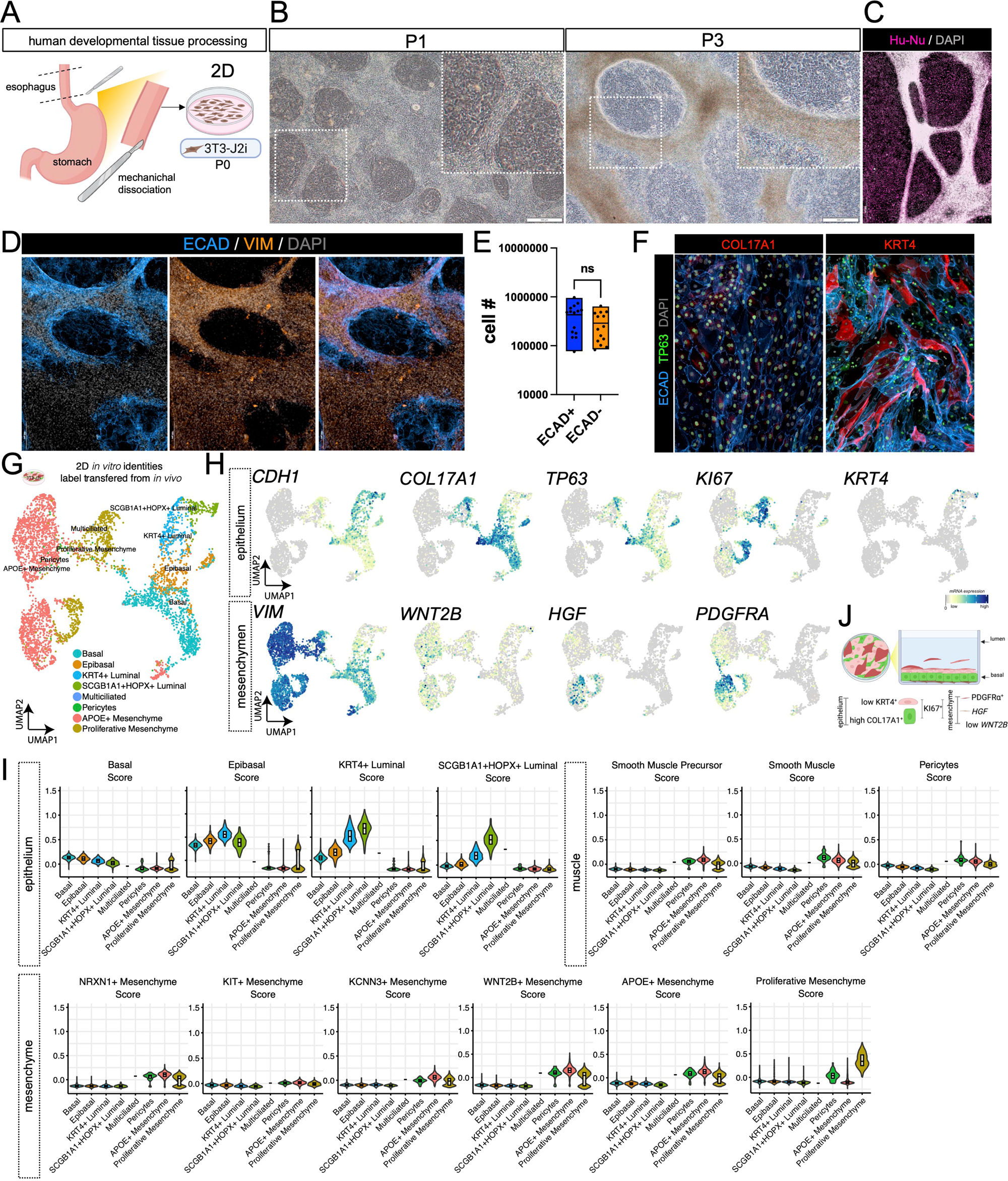
Long-Term Co-Expansion of Human Early Development Epithelial and Mesenchymal Populations *In Vitro*. (**A**) Schematic of the sample preparation workflow for *in vitro* co-expansion, including mechanical dissociation with a scalpel and co-culture with 3T3-J2i feeder cells. (**B**) Bright-field microscopy images of primary mix cells (PMC) *in vitro* cultures at passage 1 (P1) and passage 3 (P3), showing progressive expansion and illustrating the retention of distinct cell morphologies over time. (**C**) Immunofluorescence staining using human-specific nuclear antibody (Hu-Nu) to confirm the presence of human cells in the PMC *in vitro* culture. (**D**) Immunostaining for E-cadherin (ECAD) and Vimentin (VIM) to distinguish epithelial and mesenchymal populations *in vitro*. (**E**) Flow cytometry quantification of ECAD⁺ epithelial cells across multiple samples (HT n=2) and passages, showing consistent expansion of both epithelial and non-epithelial populations in co-culture. Statistical analysis indicates no significant difference (ns) or enrichment of this cell types over time. (**F**) Immunofluorescence for COL17A1, KRT4, ECAD, TP63 and DAPI confirms presence of basal and progenitor epithelial markers. **G)** Label transfer analysis mapping 2D-expanded cells onto *in vivo* epithelial clusters, showing retention of key epithelial identities from esophageal tissue. **(H)** Feature plots of key genes (*CDH1, COL17A1, TP63, KI67, KRT4, VIM, WNT2B, HGF, PDGFRA*) highlight transcriptional diversity and epithelial-mesenchymal cell retention. **(I)** Violin plots showing lineage scores grouped by predicted identity based on label transfer, including basal epithelial, epibasal, luminal, smooth muscle progenitor, smooth muscle, and pericyte populations. **(J)** Summary schematic of cell type composition in the 2D-expanded cultures.

Flow cytometry analysis across multiple samples and passages showed similar proportions of ECAD⁺ (epithelial) and ECAD⁻ (non-epithelial) cells, with no significant difference in cell numbers between these populations (Figure 3E), suggesting this method expands both epithelial and mesenchymal populations. Immunofluorescence staining confirmed the presence of key epithelial markers, including COL17A1 (basal cells), TP63 (basal/epibasal cells), KRT4 (mid-luminal cells) and ECAD, which are characteristic of basal and progenitor epithelial cells *in vivo* (Figure 1C, Figure 3F). We used passage 3 cells, plated 150,000 cells into a 6-well plate, and cultured them until day 14. To assess the transcriptional fidelity of the 2D-expanded cultures, we performed scRNA-seq. To validate the developmental relevance of the *in vitro* system, we performed label transfer analysis, mapping the 2D-expanded cells onto *in vivo* early development esophageal cells. This revealed that key epithelial identities (n=4 2D *in vitro* vs n=5 *in vivo*), including basal, epibasal populations, and luminal cells were retained 2D *in vitro* (Figure 3G). For mesenchyme populations, label transferred suggested the maintenance of pericytes, proliferative mesenchyme, and the APOE+ mesenchyme cluster (n=3 2D-*in vitro* vs n=8 *in vivo*) (Figure 3G). Feature plots of genes such as *CDH1, COL17A1, TP63, KI67, KRT4, VIM, WNT2B, HGF*, and *PDGFRA* confirmed their spatially distinct expression patterns within the 2D clusters (Figure 3H). Confidence in the label transfer was visualized using violin plots showing lineage scores grouped by predicted identity based on the label transfer, which further confirmed the presence of distinct epithelial populations, including COL17A1^+^ basal, epibasal, proliferative, KRT4⁺, and HOPX^+^ luminal as well as smooth muscle progenitor, smooth muscle, and pericyte populations (Figure 3I). This data suggests that the full mesenchymal diversity is not maintained *in vitro;* however, maintenance and expansion of some mesenchymal populations represents the first tissue derived long-term human esophagus culture model that includes mesenchymal components (Giroux, Lento et al. 2017, Kasagi, Chandramouleeswaran et al. 2018, Ferrer-Torres, Wu et al. 2022, Milne, Mustafa et al. 2024). A schematic summarizing the 2D *in vitro* system highlights the spatial organization of basal epithelial cells expressing COL17A1, proliferative cells expressing KI67, and mesenchymal cells expressing *PDGFRA* and low *WNT2B* (Figure 3J).

### Low-Adherence Suspension Culture Enables Physiologically Relevant Esophagoid Formation with Stromal-Epithelial Architecture

The 2D primary mixed culture (PMC) expansion model successfully maintains cell types from both epithelial and mesenchymal populations. However, epithelium appeared to be maintained as colonies surrounded by mesenchyme, and the epithelial-mesenchymal interactions did not represent the the spatial complexity and organization of the native esophagus (Figure 3B-D). We then wanted to determine if epithelial-mesenchymal interactions would self-organized in a 3-dimensional model. We embedded the cells into Matrigel or seeded the cells into a low adherence suspension plate (Brandenberg, Hoehnel et al. 2020) (Supplemental Figure 2A). Brightfield imaging of organoid formation in both suspension and Matrigel-based systems revealed that suspension cultures produced larger, more organized organoids over time, while Matrigel-based cultures yielded smaller and less organized structures (Supplemental Figure 2B). Quantitative analysis confirmed significantly greater growth in suspension organoids compared to Matrigel organoids across multiple time points (****p < 0.0001, Supplemental Figure 3C). Hematoxylin and eosin (H&E) staining further highlighted the structural organization of suspension organoids, which displayed well-stratified epithelial layers surrounding a central mesenchymal population, a feature less pronounced in Matrigel organoids (Supplemental Figure 2D). H&E staining of Day 14 Matrigel organoids shows only a small population of epithelial cells, with markedly reduced size compared to the fully stratified epithelium observed in suspension cultures at the same time point. (Supplemental Figure 2D). Immunofluorescence staining for ECAD (epithelial marker) and VIM (mesenchymal marker) confirmed the presence of distinct epithelial and mesenchymal compartments, with suspension organoids exhibiting more robust epithelial-mesenchymal organization when compared to Matrigel (Supplemental Figure 2D). Suspension organoids expressed the full spectrum of epithelial cell types, and formed the complete epithelial stratified layer, with expression of COL17A1 (basal progenitor expressing cell restricted to the cells near the mesenchymal population), followed by epibasal marker LY6D, and middle-to luminal differentiation marker KRT4. Interestingly, the suspension organoids also express the luminal marker CRNN, a differentiation marker not observed at this developmental time point in the matched *in vivo* sample (Supplemental Figure 2E). This suggests further differentiation of the organoids grown in suspension. Taken together, we describe this model as an esophagoid model which features a central mesenchymal core surrounded by epithelial basal stem cells and a fully stratified epithelium, recapitulating key aspects of human esophageal architecture and epithelial-mesenchymal interactions.

Given that suspension culture yielded complex epithelial-mesenchymal esophagoids, we wanted to determine if organoids could be produced at a larger scale using engineered microwells. We proceeded to optimize organoid size, uniformity, and the number of organoids formed per well by utilizing engineered agarose microwells of different diameters (500 µm to 800 µm) and comparing these to organoids formed in 96-well low adherence plate. All these systems allow for the cells to form 3D organoids without attaching to the surface. Organoids grown in smaller microwells (500 µm and 800 µm) were more uniform in size and organization, as well as in the 96 well plate (Supplemental Figure 3C), while the larger microwell (1600 µm) produced larger and more heterogeneous organoids (Supplemental Figure 3A-C). Immunofluorescence analysis revealed that all systems supported clear basal-to-luminal organization, with TP63 and COL17A1 marking basal progenitor cell types in a single layer surrounding the mesenchymal core (Supplemental Figure 3B). Quantification of organoid size over time showed that organoids in the larger microwell grew significantly larger than those in smaller microwells, but with greater variability (Supplemental Figure 3C). Nonetheless, the 96-well consistently generated on average one organoid per well, as well as more uniformity in size over time, when compared to the microwells system (Supplemental Figure C-D). These results demonstrate that the smaller engineered microwells systems as well as the 96-well low adherence plate can reliably generate self-organizing esophagoids with mesenchyme and epithelium and highlights the utility of these scalable platforms for modeling physiologically relevant epithelial-stromal interactions in both healthy and disease contexts.

Given the improved uniformity and cellular organization observed in the 96 well plate wells low adherence system, we next focused on characterizing the cellular temporal dynamics and reproducibility of this system. Brightfield imaging revealed robust organoid growth over time, with a significant increase in size between Day 6 and Day 14 (Figure 4A). Immunofluorescence staining revealed the presence of proliferative epithelial cells within the epithelial and stromal compartments, marked by KI67, alongside basal progenitors expressing TP63 and COL17A1 (Figure 4B, 4C). Differentiation markers such as KRT4 confirmed the formation of middle to luminal layers, indicating recapitulation of epithelial stratification observed in the *in vivo* esophagus (Figure 4D). Within the stromal compartment, PDGFRA expression was restricted to VIM⁺ mesenchymal cells (Figure 4E), and *HGF* was similarly confined to the mesenchymal population (Figure 4F). In contrast, *WNT2B* was expressed in both epithelial and mesenchymal compartments, suggesting a loss of regional specificity for this marker in this organoid model (Figure 4F).

**Figure 4.**
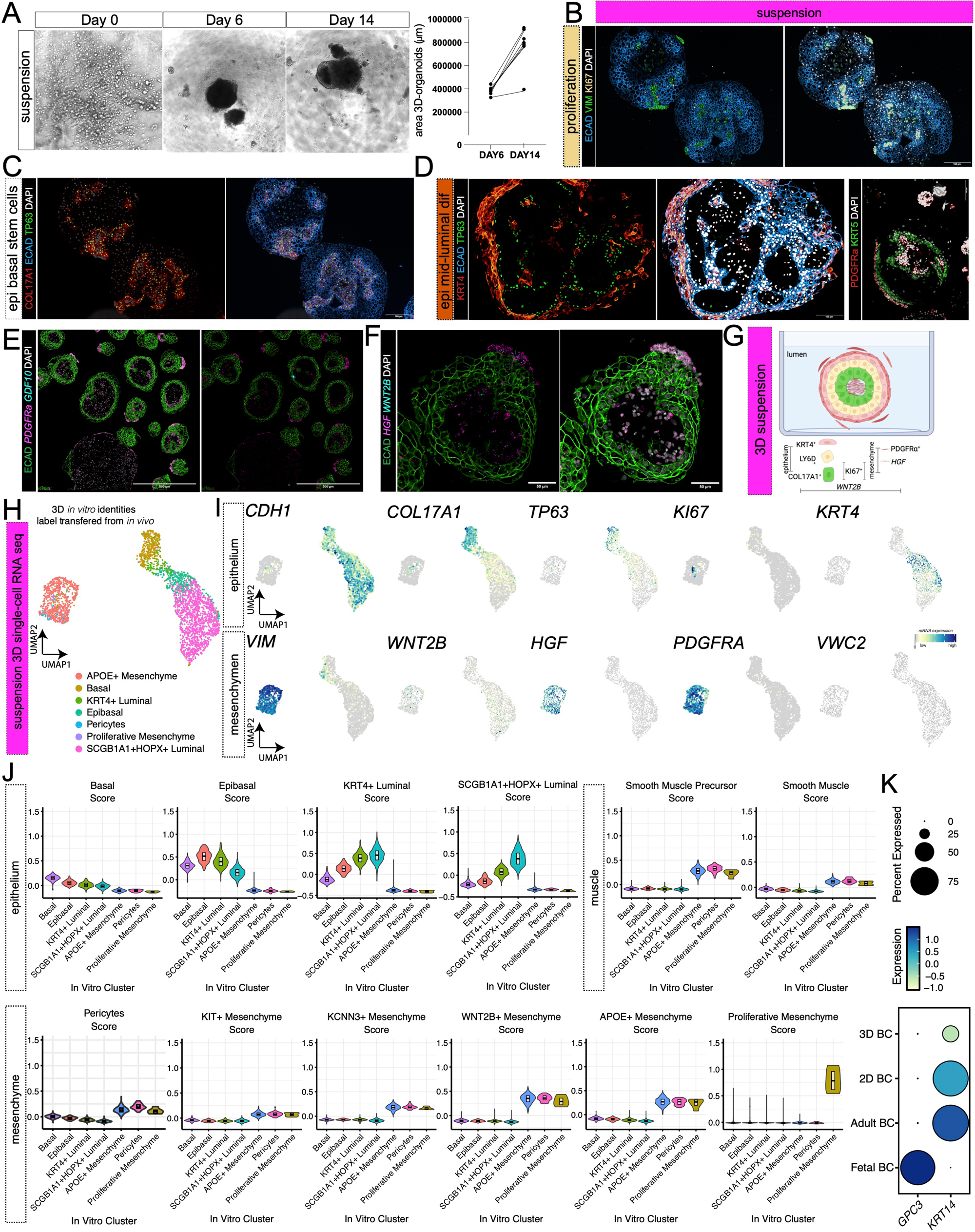
Self-Organization and Growth of Human Esophageal Cells in Suspension Culture. (**A**) Bright-field microscopy images of single-cell cultures showing self-organization into 3D organoid-like structures by day 6, with continued growth and increased size by day 14. Quantification of organoid area demonstrates significant size growth between day 6 and day 14. (**B**) Immunofluorescence staining of day 14 3D-suspension organoids reveals spatial self-organization of epithelial (E-cadherin [ECAD]+) and mesenchymal (Vimentin [VIM]+) populations, with KI67+ proliferative cells detected in both populations. (**C**) Immunofluorescence staining for COL17A1 and TP63 demonstrates the localization of epithelial basal stem cells adjacent to mesenchymal populations. (**D**) Differentiated KRT4+ epithelial cells are predominantly observed on the outer regions of the suspension organoids. (**E–F**) Immunofluorescence combined with FISH highlights the expression and spatial distribution of specific mesenchymal markers within the suspension organoids. (**G**) Schematic cross-section of a 3D organoid, labeling distinct regions and cell types including basal, luminal, and mesenchymal compartments. (**H**) Label transfer analysis mapping 3D-suspension organoids cells onto *in vivo* clusters, showing retention of key epithelial and mesenchyme identities from early development esophageal tissue. (**I**) UMAP plots of key genes for epithelium and mesenchyme clusters. (**J**) Violin plots comparing gene expression score levels across predicted cell identities and conditions (*in vivo* vs. *in vitro*), highlighting transcriptional fidelity and lineage representation. (**K**) Expression levels of GPC3 and KRT14 in adult basal cells (BC) and fetal basal cells compared with 2D and 3D basal cell.

To further validate cellular composition, we performed single-cell RNA sequencing (scRNA-seq) on passage 3, Day 14 suspension organoids. To determine the identity of the clusters, we performed 3D *in vitro* label transfer to *in vivo* identities. This analysis identified distinct epithelial (n=4 *in vitro* vs n=5 *in vivo*) and mesenchymal populations (n=3 *in vitro* vs n=8 *in vivo*), including basal, proliferative, luminal, and stromal clusters containing pericytes and APOE+ mesenchyme (Figure 4H–I). Feature plots of key markers such as *CDH1, COL17A1, TP63, KI67, PDGFRA*, and *WNT2B* confirmed the transcriptional identity distribution of these populations (Figure 4I). Finally, to assess the transcriptional fidelity of the 3D suspension-grown esophagoids, we compared gene expression profiles across predicted cell identities between *in vivo* early developmental samples and *in vitro* organoids. Violin plots revealed strong concordance in the expression of lineage markers, including those for basal, epibasal, and luminal epithelial compartments (Figure 4J). Notably, we observed a loss of the ciliated luminal component that is present *in vivo* (Figure 4J). However, as noted earlier, a CRNN positive layer is formed in these organoids (Supplemental Figure 2E), a phenotype observed in adult tissues, suggesting enhanced maturation and squamous differentiation *in vitro*. This observation prompted us to examine the expression levels of the early developmental marker *GPC3* and the adult basal progenitor marker *KRT14* in both 2D and 3D systems (Figure 4K). We found that *GPC3* expression was low in both culture systems, whereas *KRT14* expression was comparatively high (Figure 4K). These findings indicate that although early developmental human esophageal cells can be cultured and expanded *in vitro*, they undergo maturation and do not retain their early developmental identity (Figure 4K). To determine the mesenchymal populations retained in the 3D *in vitro* system, we examined transcriptional profiles across mesenchymal clusters. The violin plots show that the 3D cultures exhibit strong transcriptional similarity to many *in vivo* mesenchymal clusters, suggesting that mesenchymal populations are preserved, although their identities appear more shared and less distinct than those observed in epithelial clusters (Figure 3J). Taken together, these data confirm that suspension esophagoids maintain key transcriptional and structural features of the human esophagus. This supports their utility as a physiologically relevant model for studying epithelial stromal interactions, lineage dynamics, and disease modeling in a scalable, high throughput format.

### Epithelial-Stromal Interactions Are Required For 3D Esophagoid Formation and Maintenance

We then sought to determine if epithelial-mesenchymal interactions are important for forming esophagoids, so we purified epithelial or mesenchymal components from the 2D culture using fluorescence-activated cell sorting (FACS) for E-cadherin (ECAD) as a marker. (Figure 5A). Following sorting, both populations were successfully re-plated and expanded as pure cultures in 2D (Figure 5B). ECAD-positive cells exhibited a honeycomb-like morphology and grew in tightly packed colonies, whereas ECAD-negative stromal cells displayed elongated morphology and grew sparsely in culture. (Figure 5B). Immunofluorescence confirmed robust ECAD expression in ECAD-positive cultures and its absence in ECAD-negative culture (Figure 5C–D). Quantification revealed a statistically significant difference in ECAD expression between the two populations (unpaired t-test, *p* = 0.0014). Doubling time analysis in 2D indicates that stromal cells proliferate faster than epithelial cells, with an average doubling time of 2 days compared to 3 days for epithelial cells (p < 0.001) (Figure 5E). We next assessed whether these purified populations retained the ability to form 3D structures in suspension. ECAD-positive cells formed 3D structures significantly faster than other conditions, with an average formation time of 18.04 ± 0.75 hours (mean ± SEM), compared to 42.93 ± 2.98 hours for ECAD-negative cells, 43.57 ± 2.55 hours for primary mixed cultures (PMC), and 43.13 ± 2.27 hours for 1:1 ECAD-positive/ECAD-negative co-cultures (Figure 5F-G). Despite their rapid formation, ECAD-positive 3D structures failed to maintain structural integrity by Day 14 (Figure 5H-I). In contrast, ECAD-negative, PMC, and mixed cultures retained their 3D architecture over time (Figure 5H–I). Quantitative analysis of organoid area at Day 14 revealed that ECAD-positive cultures had significantly smaller organoids compared to ECAD-negative, PMC, and 1:1 mixed culture (Figure 5I). Immunofluorescence of Day 14 3D structures revealed that ECAD-positive cultures lacked COL17A1-positive epithelial stem cells, which are essential for epithelial regeneration. VIM staining confirmed that ECAD-negative cells were stromal in nature, lacking ECAD expression and positive for VIM. Organoids from mixed cultures that contained both epithelium and non-epithelial cells, either from PMC or 1:1 mixed culture, exhibited a layered organization with a stromal center, epithelial progenitor cells, and differentiated stratified cells (Figure 5J). All together, we show the successful purification of human esophageal epithelial and stromal components and show that the stromal cells are necessary for 3D esophagoid formation.

**Figure 5.**
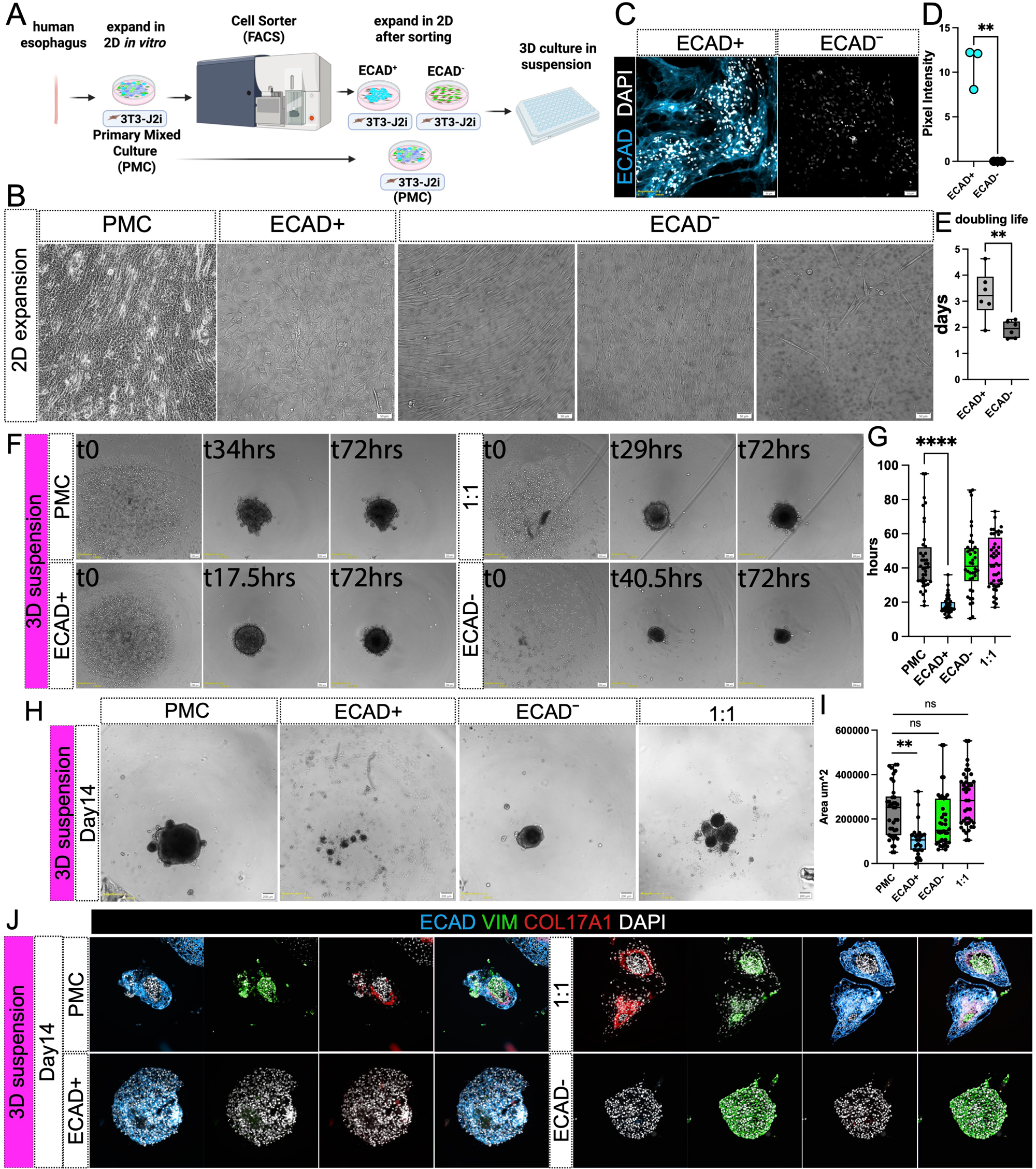
Purification and functional assessment of epithelial and stroma populations in human esophagus-derived cultures. (**A**) Schematic of the experimental workflow: primary human esophagus cells were expanded in 2D, sorted by FACS into ECAD⁺ (epithelial) and ECAD⁻ (stromal) populations, re-expanded in 2D, and cultured in 3D suspension. (**B**) Representative phase contrast images of 2D-expanded cultures. ECAD⁺ cells formed tightly packed colonies with honeycomb-like morphology, while ECAD⁻ cells exhibited elongated shapes and grew with intercellular spacing. (**C–D**) Immunofluorescence and quantification of ECAD expression confirmed successful separation of epithelial and stromal populations (p = 0.008, unpaired t-test). (**E**) Doubling time analysis in 2D indicates that stromal cells proliferate faster than epithelial cells, with an average doubling time of 2 days compared to 3 days (p < 0.001) (**F**) Time-lapse imaging of organoid formation in suspension culture. ECAD⁺ cells formed 3D structures significantly faster than ECAD⁻, PMC, or 1:1 mixed culture. (**G**) Quantification of organoid formation time across conditions. ECAD⁺ cells formed organoids at 18.04 ± 0.75 hours, significantly faster than ECAD⁻ (42.93 ± 2.98 hours), PMC (43.57 ± 2.55 hours), and 1:1 mix (43.13 ± 2.27 hours) (p < 0.001, Tukey HSD). (**H**) Representative images of Day 14 organoids. ECAD⁺ organoids showed structural collapse, while ECAD⁻, PMC, and mixed cultures retained 3D integrity. (**I**) Quantification of organoids area at Day 14. ECAD⁺ organoids were significantly smaller (102,842 ± 14,255 µm²) than ECAD⁻ (194,289 ± 19,837 µm²), PMC (236,204 ± 19,420 µm²), and 1:1 mix (286,909 ± 19,047 µm²) (F = 14.99, p < 0.0001, ANOVA; p < 0.01, Tukey HSD). (**J**) Immunofluorescence staining of Day 14 organoids for ECAD, VIM, COL17A1, and DAPI. ECAD⁺ organoids lacked COL17A1⁺ epithelial basal progenitor cells, while organoids containing stromal cells showed organized layers with stromal centers, epithelial basal progenitors, and differentiated stratified cells.

### WNT2B Signaling Regulates Esophageal Epithelial Progenitor Dynamics

We observed that the mesenchyme plays a significant role in supporting the structural organization of esophageal organoids in vitro (Figure 5H–J). We also detected retention of WNT2B expressing mesenchyme in both 2D and 3D in vitro models (Figure 3H, Figure 4I). To further investigate the role of WNT2B in esophageal epithelial development, we analyzed primary tissue samples from WNT2B loss-of-function (LOF) patients and examined the effects of WNT signaling modulation in vitro using 3D suspension organoids. Histological analysis revealed that WNT2B LOF tissues exhibited a thinner, disorganized epithelium with reduced cellularity compared to controls, as shown by H&E and Masson’s trichrome staining (MTS) (Figure 6A). Immunofluorescence staining for β-catenin, TP63, and KI67, showed an increase in proliferating TP63+ progenitors in WNT2B LOF tissues (Figure 6B, C). The increase in proliferating TP63+ progenitor cells suggests that WNT2B suppresses self-renewal of developing basal progenitor cells. In addition, papillae were significantly larger in WNT2B LOF esophageal tissues relative to healthy controls (Figure 6B-C), suggesting that diminished WNT activity alters epithelial–stromal organization and drives enhanced papillary extension.

**Figure 6.**
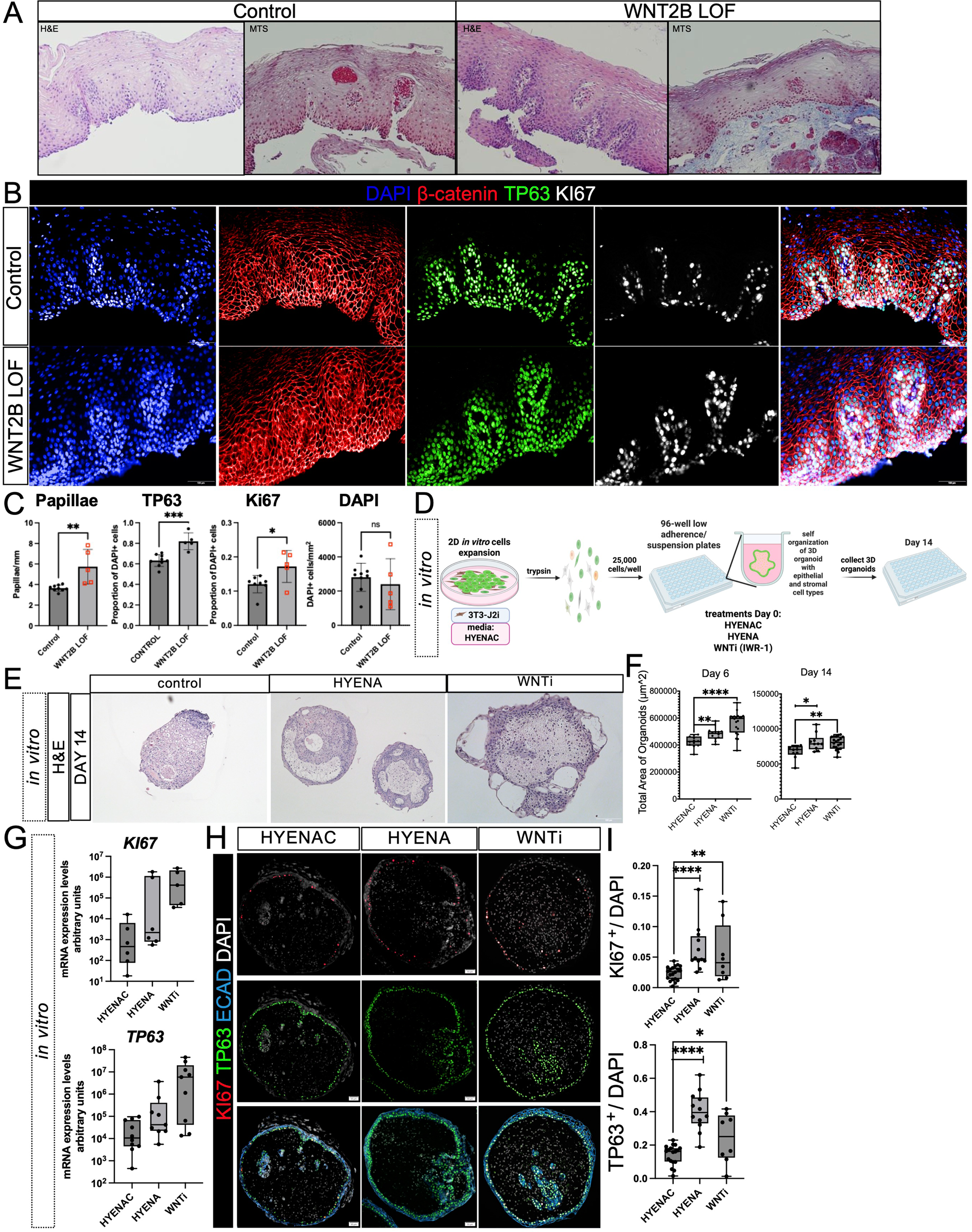
The Role of WNT2B in Esophageal Epithelial Growth and Stratification. (**A**) Histological analysis of esophageal tissues in control and WNT2B loss-of-function (LOF) models. Hematoxylin and eosin (H&E) and Masson’s trichrome staining (MTS) reveal reduced epithelial thickness and structural disorganization in WNT2B LOF tissues compared to controls. (**B**) Immunofluorescence staining for B-catenin (red), TP63 (green, basal progenitor marker), KI67 (white, proliferation marker), and DAPI (blue, nuclei) shows increased basal progenitors and proliferative cells in WNT2B LOF tissues compared to controls. Scale bar: 50 µm. (**C**) Quantification of TP63+ and KI67+ cells, DAPI+ nuclei, and epithelial papillae size (mm) in control and WNT2B LOF tissues. Bar graphs show a significant increase in TP63+ and KI67+ cells and more papillae in WNT2B LOF tissues. Data are presented as mean ± SEM, *p < 0.05, **p < 0.01. (**D**) Schematic of the experimental workflow for treating 3D esophageal organoids with WNTi-2b over 14 days, including cell seeding, drug treatment, and endpoint analyses. (**E**) H&E-stained sections of esophageal organoids under control, HYENA, and WNTi treatment conditions, showing structural differences in epithelial stratification. (**F**) Quantification of total organoid size on Days 6 and 14. WNTi-treated organoids are significantly bigger than control and HYENA-treated organoids at both time points. Data are presented as mean ± SEM, ****p < 0.0001, **p < 0.01, *p < 0.05. (**G**) mRNA quantification of Ki67 and TP63 expression across treatment conditions. (**H**) Immunofluorescence staining of 3D organoids at Day14 for KI67 (red), TP63 (green), ECAD (cyan, epithelial marker), and DAPI (nuclei). Scale bar: 50 µm. (**I**) Quantification of KI67+ and TP63+ cells in 3D organoids under control, HYENA, and WNTi conditions. WNTi treatment significantly increases the proportion of KI67+ and TP63+ cells. Data are presented as mean ± SEM, ****p < 0.0001, **p < 0.01, *p < 0.05

We then sought to understand whether the esophagoid model could be used to interrogate WNT signaling *in vitro,* and if modulating WNT signaling would mimic the *in vivo* WNT2B loss-of-function phenotype (Figure 6D). Given that 2D cultures are grown in control media which possesses the GSK3b antagonist, CHIR99021 (see methods) we carried out experiments in 3D esophagoids, which were culture in control media, media without CHIR99021, and WNT inhibitor (WNTi) (IWR-1) (Figure 6D). In 3D suspension organoids, brightfield imaging revealed robust growth under control conditions, while CHIR99021 removal or WNT inhibition resulted in even larger organoids, a statistically significant finding (Figure 6E–F). Despite increased size, H&E staining revealed that WNTi-treated organoids developed thinner, poorly organized epithelial layers, resembling the WNT2B LOF phenotype observed *in vivo* (Figure 6A, E). To assess whether WNT inhibition affected progenitor cell dynamics, we quantified KI67 and TP63 expression at both the transcription and protein levels. qPCR analysis showed a significant upregulation of *KI67* and *TP63* mRNA in WNTi-treated organoids compared to controls and CHIR99021-removal (Figure 6G). These findings were corroborated by immunofluorescence staining for KI67, TP63, which revealed an expansion of proliferative and basal progenitor populations under WNT inhibition (Figure 6H, I). These findings demonstrate that WNT plays a critical role in esophageal epithelial growth by repressing progenitor cell proliferation and restricting TP63 expression. Both *in vivo* and *in vitro* models highlight the importance of WNT signaling in maintaining epithelial homeostasis and structural integrity during esophageal development.

Importantly, these results also emphasize the physiological relevance of the esophagoid model, which successfully mimics key features of healthy versus diseased esophageal tissue. The ability of the model to recapitulate WNT2B LOF phenotypes, including disrupted epithelial architecture by WNT inhibition which lead to an increase in proliferation and basal (TP63^+^) progenitor cells, highlights its utility for studying epithelial-stromal interactions and for modeling developmental disorders and disease states of the esophagus.

## Discussion

Understanding the cellular and molecular mechanisms that regulate esophageal development is critical for advancing regenerative strategies and disease modeling. In this study, we present a multi-platform approach that integrates transcriptional profiling, spatial mapping, and *in vitro* modeling to distinguish the epithelial and mesenchymal dynamics of the human esophagus from early development to adulthood. Our comparative analysis of early developmental and adult esophageal epithelium revealed conserved lineage markers alongside stage-specific populations, including multiciliated cells and GPC3⁺ basal progenitors during early development, and KRT14⁺ basal cells together with CRNN⁺ cornified luminal cells in adults. These findings align with previous studies (Yu, Kilik et al. 2021, Ferrer-Torres, Wu et al. 2022) and provide a refined atlas of epithelial maturation, highlighting transitional states and lineage trajectories often underrepresented in adult tissue datasets. Importantly, species-specific differences emerged: whereas KRT14 is expressed early in mouse embryonic epithelium (Yu, Slack et al. 2005), we found that it is absent during early human development. Instead, GPC3 marks early human basal progenitors, while KRT14 is restricted to adult basal cells. This distinction enables the use of GPC3 and KRT14 as stage-specific markers to assess basal progenitor states in human esophageal development. In mice, KRT14 expression begins around embryonic day 15.5 (E15.5) and becomes robust during the transition from simple cuboidal epithelium to stratified squamous epithelium (E15.5–E17.5) (Yu, Slack et al. 2005), marking the establishment of the basal cell layer. In contrast, our data indicate that human esophageal development follows a distinct trajectory, with GPC3⁺ basal progenitors preceding KRT14⁺ expression, reinforcing the importance of human-specific studies for understanding epithelial lineage specification. Finally, the conserved expression of COL17A1 (basal), LY6D (epibasal), and KRT4 (middle-to-luminal) across early development and adult stages provides a robust set of markers to distinguish spatial localization and cell identities along the trajectory from basal stem cells to fully differentiated cell types in the esophageal epithelium. While most studies of basal versus differentiated states have focused on TP63 and KRT4 (Mou, Vinarsky et al. 2016, Yamamoto, Wang et al. 2016, Kasagi, Chandramouleeswaran et al. 2018, Trisno, Philo et al. 2018, Zhang, Yang et al. 2018, Ferrer-Torres, Wu et al. 2022), our analysis identifies an intermediate subpopulation between the basal zone and the onset of KRT4 expression that is marked by LY6D positivity in both adult (Ferrer-Torres, Wu et al. 2022) and early developmental samples (Figure 1). LY6D is absent in the basal progenitor layer but becomes expressed as cells exit this compartment, preceding KRT4 expression. Thus, LY6D serves as an early marker of progenitor cells transitioning out of the basal state and entering the initial steps toward luminal differentiation. This refined marker set enables more precise mapping of lineage trajectories and transitional states during esophageal development and maturation in the epithelium.

We also characterized the mesenchymal compartment of the developing esophagus, identifying transcriptionally and spatially distinct subtypes, including *WNT2B⁺*, *KIT⁺*, and *VWC2⁺* mesenchyme. These populations exhibit defined spatial relationships with the epithelium. Of importance, these findings align with the multi-omics cell census presented by Yang et al. 2025, who identified early Fibroblast Progenitors (Fib_PG) and differentiated fibroblast subtypes (Fib_1–5), Myofibroblasts (MFs), Fib_ICC, and Smooth Muscle (SM) cells. Notably, Fib_1 and MFs, which express WNT antagonists such as SFRP2, were shown to be spatially adjacent to the epithelium and critical for basal cell (BC) specification (Yang, McCullough et al. 2025). Our identification of *WNT2B⁺* and *KIT*⁺ mesenchyme matches the gene expression profile of Fib_ICC and supports a model in which mesenchymal subtypes provide localized, stage-specific signals that regulate epithelial growth and stratification. Furthermore, our observation of *VWC2*⁺ mesenchyme closely opposed to the epithelium complements their spatial mapping of Fib_1 as a signaling hub adjacent to the basement membrane. Our findings complement and validate the multi-omics cell census by Yang et al. 2025, reinforcing the concept of spatially organized stromal signaling hubs that regulate basal cell specification and epithelial stratification.

To model these interactions *in vitro*, we developed a 2D expansion system using a 3T3-J2 feeder layer (Liu, Krawczyk et al. 2017, Ferrer-Torres, Wu et al. 2022), which retained key epithelial and mesenchymal populations with transcriptional fidelity to *in vivo* counterparts. While this system allowed for expansion and characterization, it did not recapitulate esophageal spatial organization. To address this, we established a suspension-based (Brandenberg, Hoehnel et al. 2020) 3D esophagoid model that self-organizes into a physiologically relevant structure. Unlike previous systems (Andl, Mizushima et al. 2003, Kalabis, Wong et al. 2012, Giroux, Lento et al. 2017, Kasagi, Chandramouleeswaran et al. 2018, Busslinger, Weusten et al. 2021, Ferrer-Torres, Wu et al. 2022, Yang, McCullough et al. 2025), this platform self-organizes into a physiologically relevant structure with a central mesenchymal core surrounded by a stratified epithelium, recapitulating key features of *in vivo* tissue architecture. The model supports robust epithelial differentiation and captures four major trajectory cell states: COL17A1 positive basal cells, LY6D positive epibasal cells, KRT4 positive middle-to-luminal cells, and CRNN positive cornified cells, indicating enhanced maturation compared to matched developmental stages. Importantly, the esophagoid retains distinct stromal populations, including pericytes and smooth muscle cells, and demonstrates transcriptional fidelity across epithelial and mesenchymal lineages. This complexity enables functional interrogation of epithelial–stromal interactions in a context that mirrors human physiology. Furthermore, the esophagoid’s orientation allows modeling of clinically relevant insults such as reflux exposure in a manner that mimics the luminal-facing architecture of the human esophagus, an advantage not offered by other organoid systems (Giroux, Lento et al. 2017, Kasagi, Chandramouleeswaran et al. 2018, Busslinger, Weusten et al. 2021, Nakagawa, Sato et al. 2025). Finally, its scalability and reproducibility support high-throughput quantitative analysis of growth and organization, positioning this platform as a powerful tool for mechanistic studies, drug screening, and disease modeling, and circumventing scalability challenges of other systems (Lampart, Iber et al. 2023).

Functional interrogation of epithelial–stromal interactions revealed that purified epithelial cells alone were insufficient to maintain organoid integrity, emphasizing the essential role of stromal support in sustaining epithelial progenitor populations. This dependency was further validated by WNT2B loss-of-function studies, which demonstrated disrupted epithelial architecture and increased basal progenitor proliferation both *in vivo* and *in vitro*. These findings align with Yang et al. (2025), who reported that WNT inhibition (via IWP2) is required for early basal cell specification from hPSCs, whereas WNT activation supports later self-renewal, highlighting the importance of temporally regulated WNT signaling in balancing progenitor expansion and epithelial maturation (Yang, McCullough et al. 2025). Our modulation of WNT signaling in suspension organoids recapitulated these phenotypes, confirming the esophagoid model’s utility for studying disease-relevant pathways. These observations are contextualized by the broader landscape of WNT biology in the esophagus, where canonical WNT signaling has historically been poorly understood compared to its well-defined roles in other adult epithelia (Clevers, Loh et al. 2014). During early human esophageal development, suppression of WNT is required to specify esophageal progenitors from the dorsal anterior foregut (Kelley S. Harris-Johnsona 2009, Woo, Miletich et al. 2011, Trisno, Philo et al. 2018). In the adult mouse esophagus, expression of canonical WNT target genes remains low (Grommisch, Wang et al. 2024), suggesting that ongoing suppression, rather than activation, of epithelial WNT signaling is a hallmark of homeostasis. Consistent with this, somatic epithelial mutations in canonical WNT pathway genes are uncommon in adult esophageal tissues (Lin, Hao et al. 2014, Song, Li et al. 2014, Cancer Genome Atlas Research, Analysis Working Group: Asan et al. 2017, Martincorena I 2018). Together, these data support a model in which WNT signaling plays a context-dependent role in esophageal biology, requiring early inhibition for lineage specification, localized mesenchymal WNT2B input for basal progenitor regulation, and global suppression for adult homeostasis.

While this study provides valuable insights into esophageal development and epithelial-stromal interactions, several limitations should be acknowledged. First, our 3D esophagoid system lacks immune components, which are increasingly recognized as key regulators of epithelial homeostasis, wound healing, and stromal remodeling. The absence of innate and adaptive immune cells limits the physiological relevance of the model, particularly for studying inflammatory or immune-mediated esophageal diseases. Incorporating immune cells into co-culture systems or integrating organ-on-chip platforms could enhance the complexity and translational potential of future models. Second, although we demonstrate that WNT2B signaling plays a critical role in regulating basal progenitor expansion and epithelial architecture, the downstream molecular mechanisms remain unclear. Identifying the transcriptional targets and signaling cascades downstream of WNT2B, such as interactions with canonical WNT/β-catenin, non-canonical pathways, or crosstalk with BMP, TGFβ, and Hedgehog signaling, will be essential for understanding how mesenchymal cues shape epithelial fate decisions. Third, our organoid system does not fully retain the developmental stage of the tissue from which it was derived. This is evidenced by the onset CRNN expression, a marker of terminally differentiated adult luminal cells, in organoids derived from early developmental tissue, where CRNN is normally absent. This suggests that the *in vitro* environment may accelerate or alter maturation trajectories, potentially bypassing intermediate developmental states. While this may offer advantages for modeling adult-like tissue, it also highlights the need for refined culture conditions that preserve developmental fidelity when desired. Finally, it is important to note that the early developmental stage samples used for *in vitro* modeling were collected prior to the formation of submucosal glands. As a result, it remains unknown whether this model can support the expansion or maintenance of this important human cell population.

Together, our findings establish a robust framework for modeling human esophageal development and disease. The integration of single cell transcriptomics, spatial RNA and protein profiling, and scalable organoid technologies provides a powerful platform for interrogating epithelial-stromal interactions, testing regenerative strategies, and exploring the molecular underpinnings of congenital and acquired esophageal disorders.

## Conclusion

In summary, our study provides a comparative analysis and structural atlas of human esophageal development, offering novel insights into the cellular and molecular mechanisms governing epithelial and mesenchymal interactions. We established a long-term expansion method that preserves both epithelial and mesenchymal populations *in vitro*, enabling extended study of esophageal cell dynamics. Additionally, we developed a high-throughput 96-well suspension culture system that facilitates the formation of physiologically relevant esophageal organoids, faithfully recapitulating the structural and cellular complexity of the human esophagus *in vitro*. Furthermore, our findings suggest that WNT2B plays a critical role in epithelial-mesenchymal crosstalk during organ formation, both *in vivo* and *in vitro*, highlighting its potential as a key regulatory factor in esophageal development. These advancements not only deepen our understanding of esophageal biology but also establish a robust foundation for disease modeling, tissue regeneration, and therapeutic interventions.

## Star Methods

### EXPERIMENTAL MODEL AND STUDY PARTICIPANT DETAILS

#### Ethics Statement on the Use of Human Fetal Tissue and Human Subjects

This study utilized human fetal esophageal tissue obtained through ethically approved protocols and in full compliance with federal, state, and institutional regulations governing the use of human fetal tissue in research. All tissue samples were procured from established tissue banks under Institutional Review Board (IRB)-approved protocols (IRB #380), with informed consent obtained from all donors prior to tissue collection. Consent was obtained only after donors had independently consented, and no financial incentives were provided for tissue donation.

The University of Washington Birth Defects Research Laboratory (BDRL) facilitated tissue procurement, ensuring that only tissue not required for clinical care was collected. All personal identifiers were removed, and samples were coded to maintain donor anonymity. A Certificate of Confidentiality from the National Institute of Child Health and Human Development was obtained to further protect donor privacy.

The protocols and use of human fetal tissue for this research have been reviewed and approved by the Scientific Ethics Committee at the University of Colorado Anschutz Medical Campus, ensuring that all procedures meet the highest standards of ethical and scientific integrity. This research aims to advance understanding of congenital esophageal disorders by modeling early human esophageal development using organoids derived from fetal tissue. The ethical framework guiding this work emphasizes transparency, accountability, and respect for human dignity.

#### 3T3-J2 Cell Culture and Maintenance

The 3T3-J2 fibroblast feeder cell line was used to support the 2D expansion of human early development esophageal epithelial cells, and was generated per previously described (Liu, Krawczyk et al. 2017) In short, the J2 line was purchased from Kerafast and the sex was not disclosed. J2 media were prepared by combining 500 mL DMEM (REF 11960-044; no L-glutamine), 50 mL iron-supplemented bovine calf serum (ATCC 30-2030; not gamma-irradiated or heat-inactivated), 5.5 mL Pen/Strep (Gibco), and 5.5 mL L-GlutaMax (Gibco 25030-081). For thawing, frozen J2 cells (approximately 600,000 cells per vial) were quickly thawed in a 37°C water bath for 30 seconds and added to 50 mL pre-warmed media without centrifugation. Cells were seeded into T150 flasks and incubated at 37°C with 10% CO₂. Media were changed 24 hours after seeding and then every two days. Cells were cultured to 70–90% confluence before passaging. Once at the right confluency, media was aspirated, and cells were rinsed with 5 mL DPBS. A total of 5 mL of 0.05% Trypsin/EDTA was added to detach the cells. Trypsin activity was quenched by adding 5 mL of media, and cells were centrifuged at 500×g for 5 minutes. The cell pellet was resuspended in 7 mL media and triturated 20 times before being split at ratios of 1:2 or 1:6 depending on growth needs. Cells were not allowed to grow beyond 80% confluence, as over-confluence affects irradiation sensitivity. For feeder cell preparation, 3T3-J2 cells were irradiated with 30 Gy and resuspended at 1.2 × 10⁶ cells/mL in media supplemented with 5% DMSO. Aliquots of 600,000 cells in 500 µL were frozen in vials. Irradiation success was confirmed by plating 100,000 cells per well and ensuring no growth after incubation.

#### HYENAC Media Preparation for Esophageal Organoid Culture

The HYENAC medium was prepared as previously described by Ferrer-Torres et al. (2022).

#### Fluorescence In Situ Hybridization (FISH)

To visualize the spatial distribution of specific mRNA transcripts in early development esophageal tissues and organoids, fluorescence in situ hybridization (FISH) was performed using the RNAscope protocol (ACD Bio-Techne). Tissues and organoids were fixed in 4% paraformaldehyde overnight at 4°C and then embedded in paraffin. Paraffin sections were deparaffinized in Histoclear II and rehydrated through graded ethanol solutions, while cryosections were briefly washed with PBS. To enhance probe accessibility, sections underwent hydrogen peroxide treatment to block endogenous peroxidase activity, followed by antigen retrieval using a citrate-based buffer and protease digestion. For probe hybridization, target-specific probes for *WNT2B, PDGFRA, TP63*, or other markers of interest were applied, and the sections were incubated at 40°C in a humidified chamber for 2 hours. Hybridized probes were subsequently amplified using a proprietary amplification system and labeled with fluorophores. Slides were mounted using FluorSave Reagent (Milliepore-Sigma: 345789). Nuclei were counterstained with DAPI for visualization with confocal microscopy (Nikon) to capture the spatial distribution of target transcripts. Fluorescence signals and co-localization with protein markers were analyzed to confirm spatial relationships between epithelial and mesenchymal cell populations.

#### Immunofluorescence and Histological Analyses

Immunofluorescence staining of paraffin embedded tissues. Slides were deparaffinized in Histoclear II for 5 minutes (two changes) and rehydrated through graded ethanol solutions (100%, 95%, 70%, and 30% ethanol, two changes each for 3 minutes). After rehydration, slides were rinsed in double-distilled water (ddH₂O) for 5 minutes (two changes). Antigen retrieval was performed by placing the slides in sodium citrate buffer (25 mL of 10× sodium citrate buffer diluted with 225 mL ddH₂O), which was heated in a microwave for 2 minutes until cloudy. Slides were then steamed in a Tissue-Tek container for 20 minutes and allowed to cool for 15–20 minutes before being rinsed in ddH₂O for 5 minutes (two changes). After removing excess liquid, sample areas were circled with a hydrophobic pap pen, and a blocking solution was applied to prevent nonspecific binding. Slides were incubated at room temperature for 30 minutes to 1 hour in a humidified hydro-chamber. Primary antibodies, diluted in blocking solution (stocks were made up of 4.7mls of TBS, 50uls of 10% Triton X, and 250uls of Donkey serum), were applied, and slides were incubated overnight at 4°C in the hydro-chamber.

The following day, slides were washed in 1× PBS for 5 minutes (three changes) to remove unbound primary antibodies. Secondary antibodies and DAPI, diluted in blocking solution, were applied, and the slides were incubated at room temperature for 2 hours in a light-tight hydro-chamber. After final washes in 1× PBS (5 minutes, three changes), slides were mounted using ProLong Gold. Cover slipped slides were stored in the dark at 4°C until imaging. For immunofluorescence (IF) staining, the following primary antibodies were used at a 1:1000 dilution unless otherwise specified. Rabbit anti-CAV1 (HPA049326, Sigma), rabbit anti-CAV2 (HPA044810, Sigma), rabbit anti-COL17A1 (HPA043673, Sigma), goat anti-CRNN (AF3607, R&D), mouse anti-ECAD (610181, BD Biosciences), rat anti-ECAD (13-1900, Thermo Fisher), mouse anti-KI67 (652402, BioLegend), rabbit anti-KRT4 (HPA034881, Atlas), rabbit anti-LY6D (HPA024755, Atlas), goat anti-TP63 (BAF1916, R&D), goat anti-VIM (sc-7558, Santa Cruz), and mouse anti-Hu-Nu (ab191181, Abcam).

#### Preparation of pHEMA-Coated Plates for 3D Suspension Organoid Culture

To generate 3D suspension organoids, we followed previous protocols (Choi, Vodyanik et al. 2011, Capeling, Huang et al. 2022) where tissue culture plates were coated with a 10% poly(2-hydroxyethyl methacrylate) (pHEMA) solution. For the coating solution, 4 g of pHEMA (Sigma, P3932) were added to 40 mL of 95% ethanol containing 10 mM sodium hydroxide (NaOH; Fisher Scientific, S318). The NaOH acts as a solubilizing agent, but caution was exercised due to its corrosive nature, using rubber gloves and protective goggles during preparation. The solution was dissolved completely by rotating continuously at room temperature or 37°C overnight. To prevent pHEMA precipitation, the solution was shaken immediately after adding pHEMA. Once dissolved, the solution was stored at room temperature until use. In a tissue culture hood, the pHEMA solution was applied generously to cover the entire surface of the plate. Excess solution was carefully removed and returned to the primary stock, as it could be reused multiple times. The coated plates were left uncapped under ultraviolet (UV) light in the tissue culture hood overnight to ensure sterility. The following day, plates were washed once with 1× PBS before seeding cells for 3D suspension culture.

#### 3D Suspension Organoid Culture, Matrigel and engineer microwells

Dissociated esophageal cells were cultured in suspension system (Brandenberg, Hoehnel et al. 2020) or by embedding in Matrigel (Discovery Labware, Cat No. 354234), suspension culture without an extracellular matrix, or in engineered microwells. Cells were seeded in 96-well plates at densities optimized for organoid formation, (150,000 cells per well in all system). Engineered microwells made of agarose were generated as previously described (Eiken, Levine et al. 2025). Briefly, poly(dimethylsiloxane) (PDMS) (DOW Chemicals Corporation) microwell molds were generated from the surface of EZSPHERE plates (Nacalai USA) of various sizes: 500/200 μm (width/depth), 800/300 μm (width/depth), and 1600/400 μm (width/depth)(Lo(Loebel, Saleh et al. 2022). Agarose (Millipore Sigma) was dissolved at 2.2 wt% in PBS and heated until fully dissolved, 2 mLs were added to a 6-well plate, and the PDMS mold was applied. The agarose was crosslinked by cooling and the mold removed. The hydrogel microwell surface was washed with PBS and UV-sterilized before adding cells. Brightfield and immunofluorescence imaging were used to evaluate organoid structure and marker expression.

#### Flow Cytometry for Sorting ECAD+ Cells

Flow cytometry was performed to analyze cell surface markers in esophageal epithelial and mesenchymal populations. Cells were collected and dissociated into single-cell suspensions using TrypLE at 37°C until fully dissociated. An equal volume of culture medium was added to neutralize TrypLE activity, and the suspension was filtered through a 70 µm strainer to remove cell aggregates. Cells were harvested by centrifugation at 300 × g for 5 minutes at 4°C, the supernatant was removed, and the pellet was resuspended in ice-cold FACS buffer (PBS supplemented with 3% BSA). After washing twice by centrifugation at 300 × g for 5 minutes at 4°C, cells were resuspended in 100 µL of FACS buffer. For antibody staining, 0.1–10 µg/mL of a primary labeled antibody was added to each sample, with dilutions made in FACS buffer if necessary. Samples were incubated for 30 minutes at 4°C in the dark. After incubation, cells were washed twice with ice-cold FACS buffer by centrifugation and resuspended in 200 µL to 1 mL of buffer. DAPI (1 µg/mL) was added to assess cell viability. Samples were kept on ice in the dark or stored at 4°C until analysis. Flow cytometry was performed immediately for optimal results. Data were collected using a flow cytometer and analyzed using FlowJo software. Sorted ECAD+ and ECAD– cells were collected in FBS-coated tubes containing cold HYENAC medium to maintain viability. Sorted cells were then cultured with 3T3-J2i and expanded for further experiments.

#### WNT Modulation Experiments

To assess the role of WNT signaling inhibitor (IWR-1, ApexBio, Cat No. B2306 at 5uM) was added to organoid cultures. Organoid growth and morphology were monitored over 14 days. Brightfield and immunofluorescence imaging were used to evaluate organoid structure and marker expression.

#### RNA Extraction, Reverse Transcription to make cDNA, and qPCR

At day 14, we extracted the RNA from esophageal organoids using a TRIzol (Invitrogen™ TRIzol™ Reagent, Cat No. 15-596-026) and chloroform (Sigma-Aldrich, Cat No. 366927) protocol. Organoids were extracted from a 96 well plate and then spun down to remove media. 200ul of TRIzol was added, then after a brief incubation 200ul of chloroform was added, then centrifuged for 15 minutes at 12,000g. The upper clear aqueous phase containing the RNA was removed to a new tube, then isopropanol was added, then we centrifuged the tubes again for 15 minutes at 12,000g. The isopropanol was removed, and 70% ethanol was added, we centrifuged for 5 minutes at 7,500g. Ethanol was removed, and then RNA pellet was allowed to dry in a laminar flow hood. RNA pellets were resuspended in 10ul of nuclease free water, placed on a heat block preheated to 60°C then concentration was quantified using a Nanodrop.

After quantification, the amount of RNA solution containing 500ng of RNA was calculated and then added to PCR tubes, along with 4ul of iScript Supermix (Bio-Rad, iScript™ Reverse Transcription Supermix, Cat No. 1708841) and 13-15ul of nuclease free water to make a total of 20ul per tube.

After cDNA was reverse transcribed, it was stored on ice and primer master mix was created for each primer being used. The primer master mix was made up of 5ul of SYBR green (Abclonal Technologies, 2X Universal SYBR Green Fast qPCR Mix, Cat No. RK21203), 2ul of nuclease free water, and 1ul of working stock primer (per well on PCR plate). Primers were custom made by Thermo Fisher Scientific and were dry, we resuspended them to be at a concentration of 100uM. Our working stocks of primer were made using 180ul of nuclease free water, 5ul of the forward primer, and 5ul of the rear primer.

Each well received 2ul of cDNA and 8ul of master mix. PCR plate was spun down, and then amplification was quantified using a Quantstudio3 96-well rtPCR reader.

#### Single-Cell RNA Sequencing and Data Analysis

##### Preprocessing and Quality Control

Reads were mapped to human genome (GRCh38-2020-A) and gene expression matrices generated using CellRanger v7. Cell calling was also performed in CellRanger. Filtered cell by gene count matrices were imported into R v4.0 in Rstudio v1.4 and further processed using Seurat v4 – v5.

For tissue specimens, only nuclei within the following thresholds were analyzed: between 1500 and 7500 features and with less than 20% mitochondrial RNA reads. Data from developing esophagus 2D cultures and esophagoid 3D cultures was filtered to cells with between 1500 and 10000 features, with less than 20% mitochondrial reads and more than 10000 unique molecular identifiers. Developing esophagus data were also processed without filtering to recover a cluster of multiciliated cells undergoing luminal shedding. This cluster was appended to the merged developing esophagus dataset after quality control filtering and prior to data integration. *Data integration (batch-correction), dimensional reduction and initial clustering*

Each sample was normalized using Seurat::NormalizeData() and variable features were determined using Seurat::FindVariableFeatures(). Features were selected for integration between datasets from similar specimens (i.e. adult esophagus, developing esophagus, two- and three-dimensional esophagus cultures) using Seurat::SelectIntegrationFeatures(). These features were then scaled in each dataset using Seurat::ScaleData() and scaled data was used as input for Seurat::RunPCA().

The first thirty principal components (PCs) were used as input for Seurat::FindIntegrationAnchors() using the reciprocal PCA methodology, and a batch-corrected assay was generated using Seurat::IntegrateData(), again using the first thirty PCs. Seurat:CellCycleScoring() was performed on the batch-corrected assay, followed by Seurat::ScaleData() with regression of S-phase and G2/M-phase scores. The scaled, batch-corrected data was used as input for Seurat::RunPCA(). Seurat::RunUMAP() was then performed on each batch corrected dataset using the following PCs: adult esophagus: 1 to 12, developing esophagus, three dimensional esophagoid and two dimensional esophagus cultures: 1 to 14. The same number of PCs were used for Seurat::FindNeighbors(). Louvain clustering was then performed using Seurat::FindClusters at the following resolutions: adult esophagus: 0.15, developing esophagus: 0.4, 2D cultures and esophagoid 3D cultures: 0.3.

##### Subclustering

Epithelial clusters in the adult and developing esophagus data were identified on the basis of *CDH1* expression, and mesenchymal clusters were identified on the basis of *VIM* expression. lls belonging to epithelial or mesenchymal clusters were extracted into separate objects, and the integrated assay from the full dataset was used to compute new PCs using Seurat::RunPCA() for each object. Seurat::RunUMAP was then performed using the following PCs: adult esophagus epithelium: 1 to 12, developing esophagus epithelium: 1 to 6, developing esophagus mesenchyme: 1 to 10. Louvain clustering was performed using Seurat::FindClusters at the following resolutions: adult esophagus epithelium: 0.15, developing esophagus epithelium and mesenchyme: 0.4.

##### Cell type annotations

Clusters in adult and developing esophagus datasets were annotated manually based on known markers defining stratified epithelial populations in the adult esophagus (Ferrer-Torres, Wu et al. 2022) and known markers of distinct mesenchymal subtypes. When a cell type’s gene enrichment profiles did not correspond to a known cell type clusters were named to include a highly specific marker and their major cell class (i.e. *KCNN3*+ mesenchyme).

##### Label Transfer

To align developing esophagus epithelial cell type identities to adult esophagus epithelial cell type identities label transfer was performed using the uncorrected RNA assay of both datasets and the first 30 PCs. Anchors were determined using Seurat::FindTransferAnchors() and labels were transferred using Seurat::TransferData().

To infer *in vitro* cell identities from *in vivo* reference data label transfer was performed using the batch-corrected assay in reference and query datasets to generate anchors by FindTransferAnchors(k.filter = 200), and labels were transferred using Seurat::MapQuery().

##### Gene module scoring

Gene set modules for each cell type in the developing esophagus data were selected by taking the top 100 enriched genes by log-fold change as determined by Seurat::FindMarkers(). All cells from three-dimensional esophagoid and two-dimensional esophagus cell cultures were scored for all gene modules using a gene set method developed by Tirosh et al.(Tirosh, Izar et al. 2016),and implemented using Seurat::AddModuleScore().

### QUANTIFICATION AND STATISTICAL ANALYSIS

Statistical analysis of gene expression differences between cell cultures in scRNA-seq data was performed in Seurat using FindMarkers() with default settings.Quantitative analyses of organoid size, marker expression, and cell proliferation were performed using Olympus Cellsens, using the manual threshold to quantify the size of organoids in um^2, and using the adaptive threshold to distinguish marker expression from background fluorescence. Quantification of nuclei was performed using a neural network built into Cellsens which counted nuclei stained with Dapi. Statistical analyses were conducted using GraphPad Prism. Data were expressed as mean ± SEM, and significance was determined using Student’s t-tests, one-way ANOVA, or two-way ANOVA with post-hoc corrections. P-values < 0.05 were considered statistically significant.

To quantify the rate of organoid formation, we classified the organoids as fully formed when they stopped contracting. Thus, we recorded the time frame at which they stopped contracting. To double check our assessment and decrease subjectivity, we compared the size of the organoid at our recorded data point to the size of the organoid at the final recorded frame, as shown in Figure 5.

Data were tested for normality and analyzed using parametric tests appropriate for group comparisons. To compare ECAD expression levels between ECAD⁺ and ECAD⁻ populations (Figure 5D), an unpaired two-tailed Student’s t-test was used. This test was chosen because it is appropriate for comparing the means of two independent groups with continuous data.

For comparisons involving more than two groups, including organoids formation time (Figure 5F) and organoid area at Day 14 (Figure 5H), a one-way analysis of variance (ANOVA) was used to assess overall differences among the four conditions: ECAD⁺, ECAD⁻, primary mixed culture (PMC), and 1:1 ECAD⁺/ECAD⁻ mix. ANOVA is suitable for evaluating differences in means across multiple groups when the data are approximately normally distributed and variances are comparable.

Following significant ANOVA results, Tukey’s Honestly Significant Difference (HSD) post hoc test was applied to determine pairwise differences between conditions while controlling for family-wise error rate. Tukey’s HSD is appropriate for multiple comparisons and provides adjusted p-values to identify statistically significant group differences. Significance thresholds were set at p < 0.05, with exact p-values reported where applicable. Data are presented as mean ± standard error of the mean (SEM) unless otherwise noted.

#### Rigor and Reproducibility

All experiments were conducted with a minimum of two to three technical replicates per condition, and biological replicates ranged from two to four independent samples, ensuring rigor and reproducibility across assays.

## Data Availability

Fully processed and annotated RObjects for all scRNA-seq datasets analyzed in this manuscript are available at Zenodo (https://zenodo.org/uploads/17410413). Raw scRNA-seq data from *in vitro* esophagus cultures will be available on ArrayExpress at the time of pulication. Raw scRNA-seq data from adult-stage esophagus specimens is available at ArrayExpress (E-MTAB-12266). Raw scRNA-seq data from developing esophagus specimens is available at ArrayExpress (E-MTAB-10187).

## Code Availability

Code used for analysis of single-cell RNA-seq data is available from https://github.com/jason-spence-lab/Frum-et-al.-2025b.git.

## Schematics and diagrams

Schematics were created with BioRender.com and Adobe Illustrator.

## Declaration of generative AI and AI-assisted technologies in the manuscript preparation process

During the preparation of this work the author(s) used Microsoft Copilot to verify for grammatical errors. After using this tool/service, the author(s) reviewed and edited the content as needed and take(s) full responsibility for the content of the published article.

**KEY RESOURCES TABLE:**

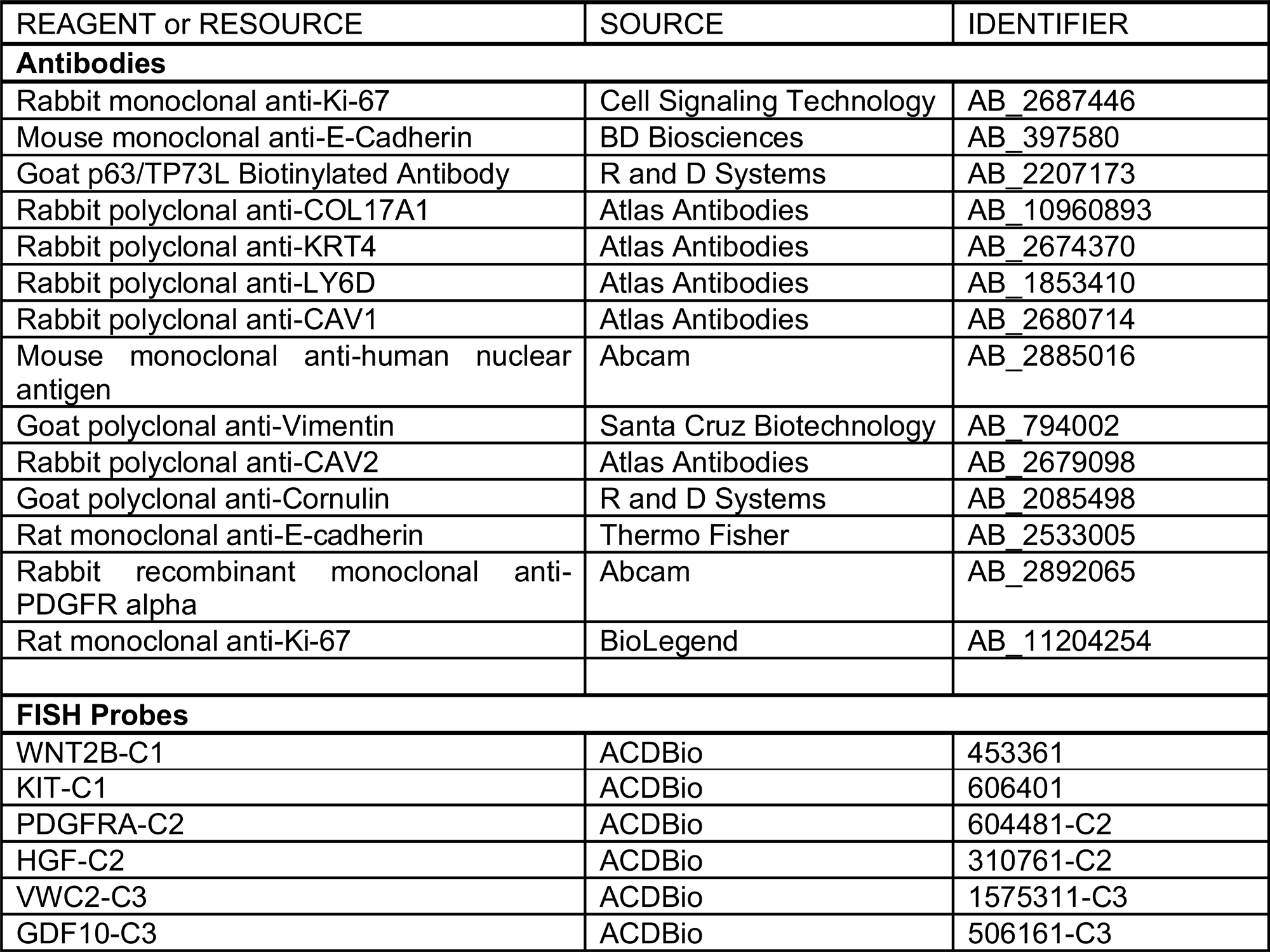
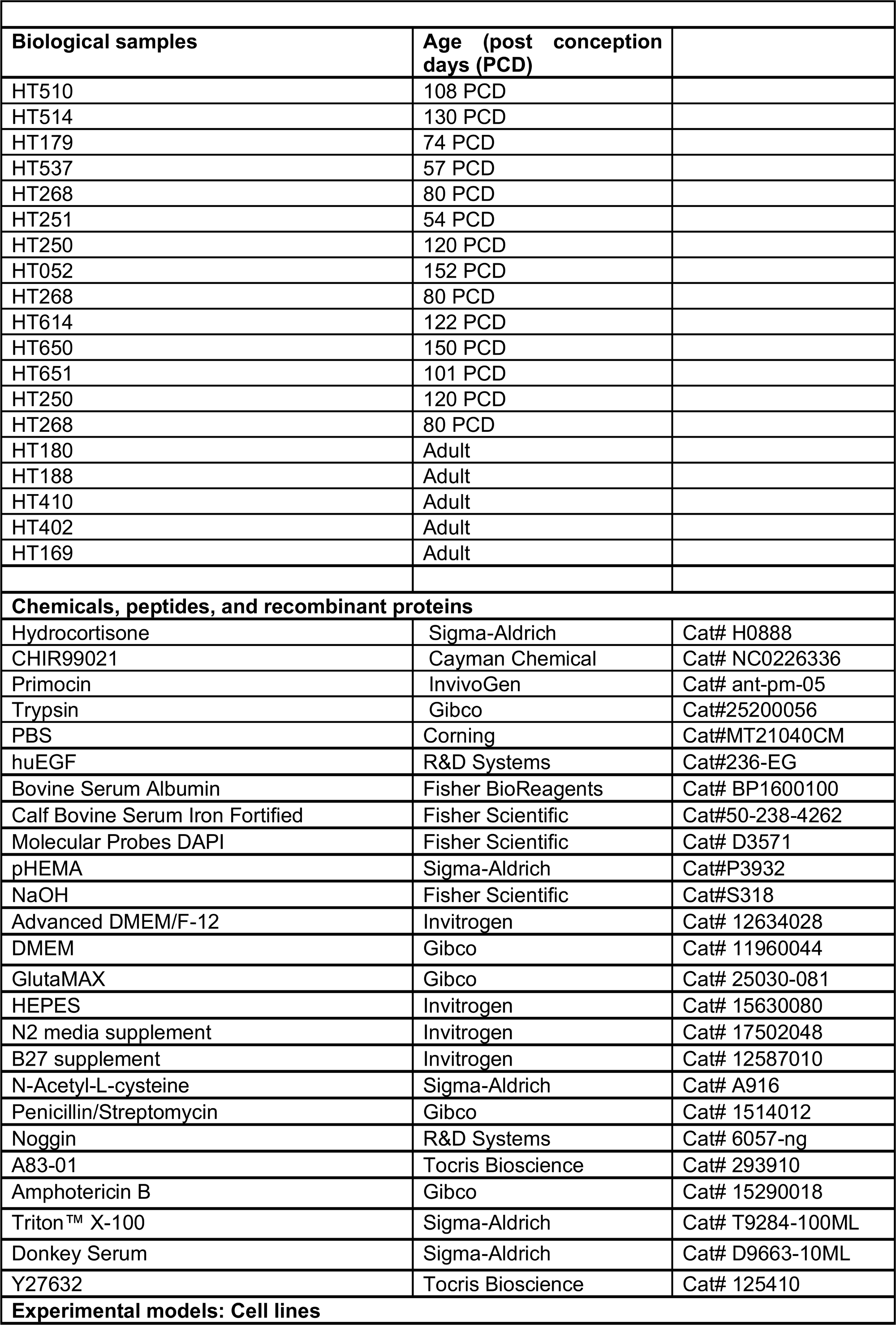
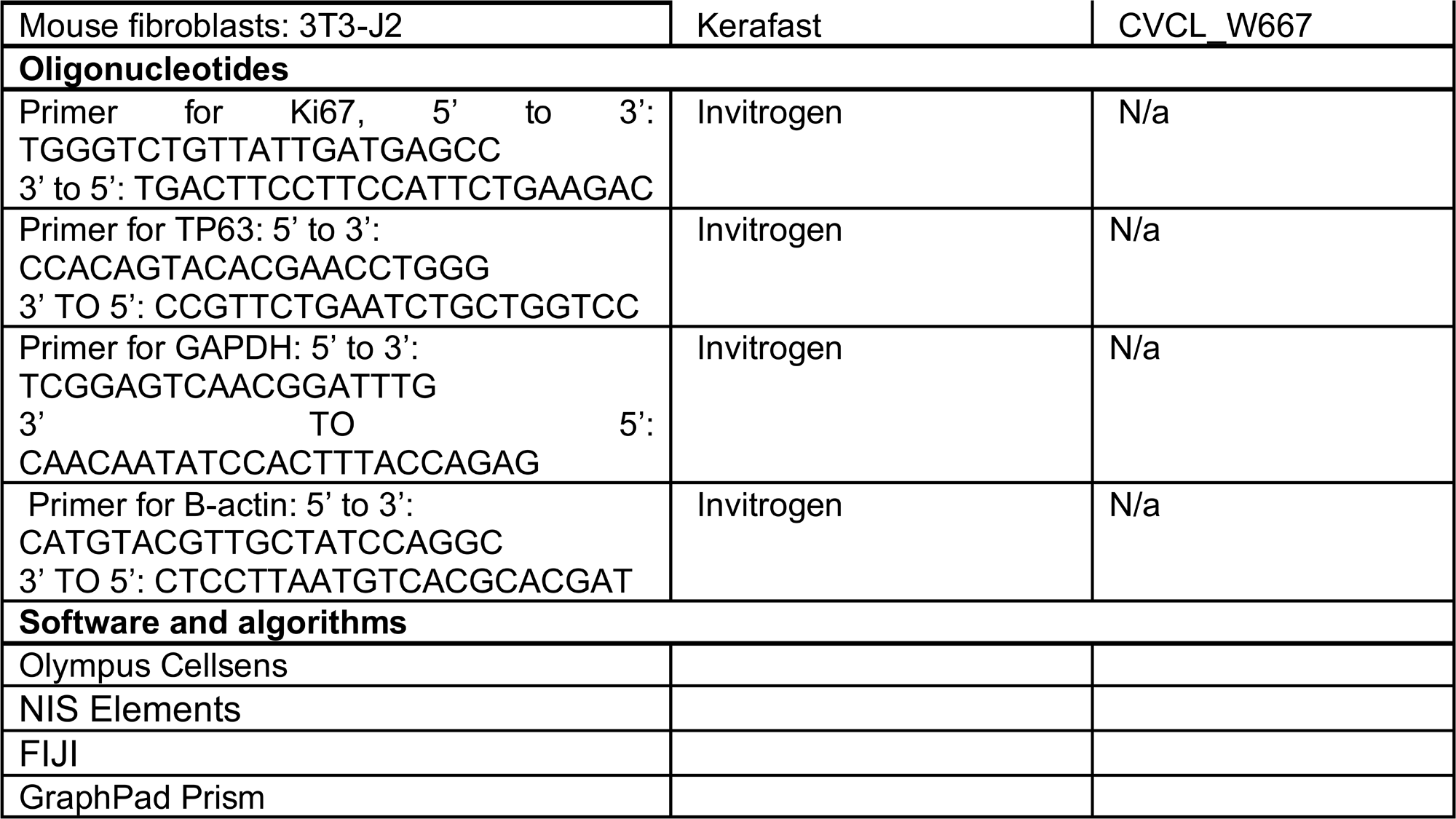

## Acknowledgements

We extend our gratitude to the University of Michigan Advanced Genomics Core for their expertise in operating the 10x Chromium single-cell capture platform and sequencing. We also thank the University of Michigan Microscopy Core for providing access to confocal microscopes and image analysis software, as well as the staff of the University of Washington Laboratory of Developmental Biology.

We appreciate the support of the University of Michigan Histology Core in processing our histology samples, with a special acknowledgment to Emma Snyder-White. We also thank the Michigan Medicine Translational Tissue Modeling Laboratory (TTML) for supplying growth medium and offering technical guidance for human *in vitro* modeling. The Translational Tissue Modeling Laboratory is a University of Michigan-funded initiative supported by the Center for Gastrointestinal Research, the Office of the Dean, the Comprehensive Cancer Center, and the Departments of Pathology, Pharmacology, and Internal Medicine, with additional support from the Endowment for Basic Sciences.

---

**Supplemental Figure 1:**
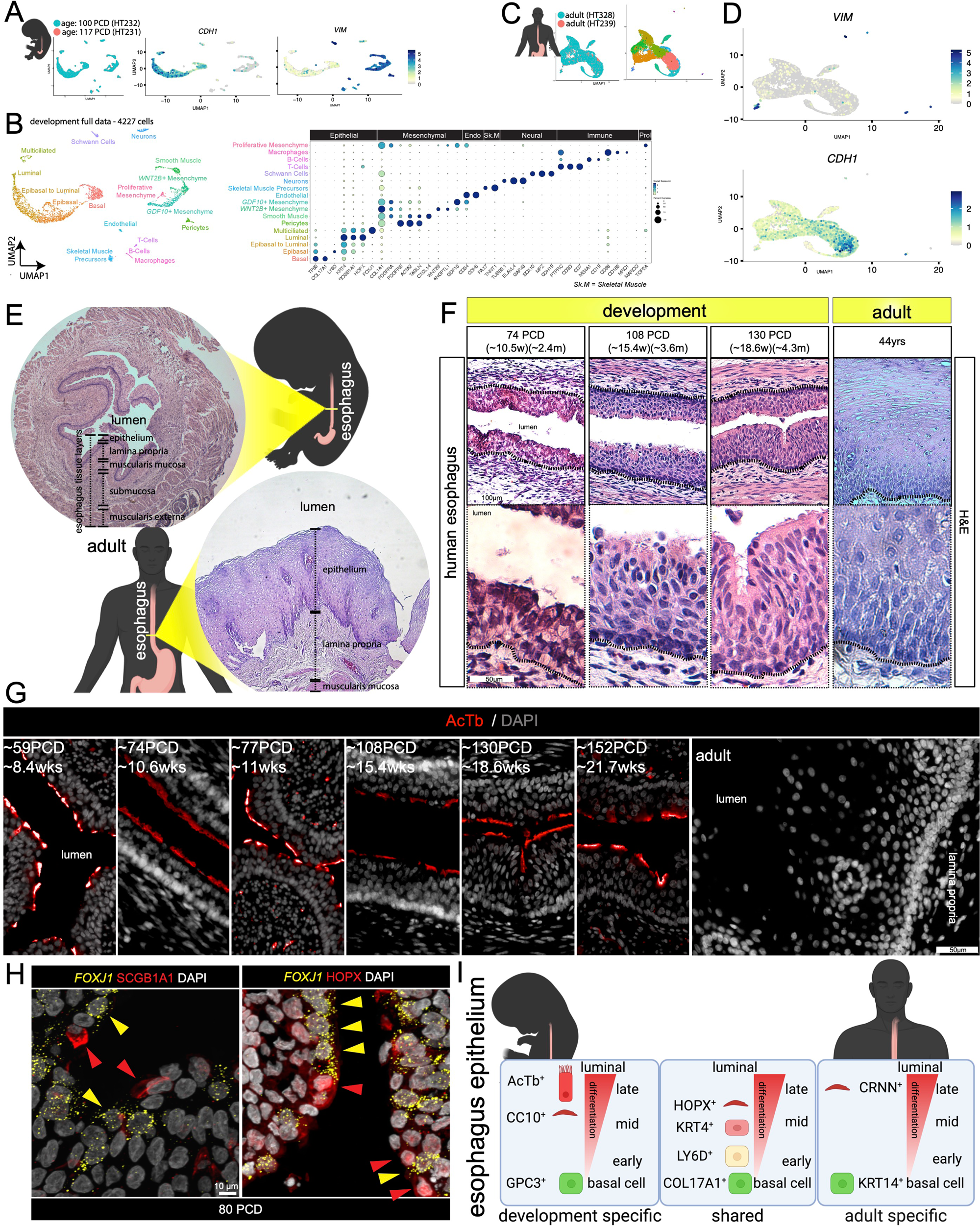
Transcriptional and Histological Characterization of Human Early Development and Adult Esophagus. (**A**) UMAP plots displaying single-cell RNA sequencing (scRNA-seq) data of early development esophageal cells (100 post-conception days [PCD] and 117 PCD) showing the expression of epithelial marker *CDH1* and mesenchymal marker *VIM*, highlighting distinct epithelial and mesenchymal populations. (**B**) Dot plot representing the scaled expression of key markers across epithelial, mesenchymal, skeletal muscle, neural, and immune cell populations, revealing transcriptional heterogeneity in the developing human esophagus. (**C-D**) UMAP plots of adult esophageal cells with feature plots showing the expression of epithelial marker *CDH1* and mesenchymal marker *VIM*. (**E**) Schematic and histological comparison of early development and adult esophageal tissue architecture. H&E staining demonstrates differences in epithelial stratification, mesenchymal organization, and muscularis mucosa between early development and adult esophagus. (**F**) H&E staining of human esophageal tissues at different developmental stages (74 days, 108 days, 130 days post-conception) and an adult esophagus (44 years). The early development epithelium undergoes progressive stratification and differentiation over time. Scale bars: 50 µm. (**G**) Immunofluorescence staining for AcTβ (red) and DAPI (nuclei) across various early development stages (59–152 PCD) and adult esophagus, showing the dynamic distribution of acetylated tubulin in the human esophagus epithelium. (**H**) Immunofluorescence staining for SCGB1A1 (red, left) and HOPX (red, right) coupled with FISH of *FOXJ1* in 80-day early developmental esophageal tissue. Scale bar: 10 µm. (**I**) Schematic summarizing epithelial populations in early development and adult esophagus. Early development-specific markers (AcTβ+, CC10+, GPC3+) distinguish luminal, mid, and early basal cells, whereas adult basal cells express KRT14 and luminal cells express CRNN. Shared markers include HOPX, KRT4, LY6D, TP63, and COL17A1.

**Supplemental Figure 2:**
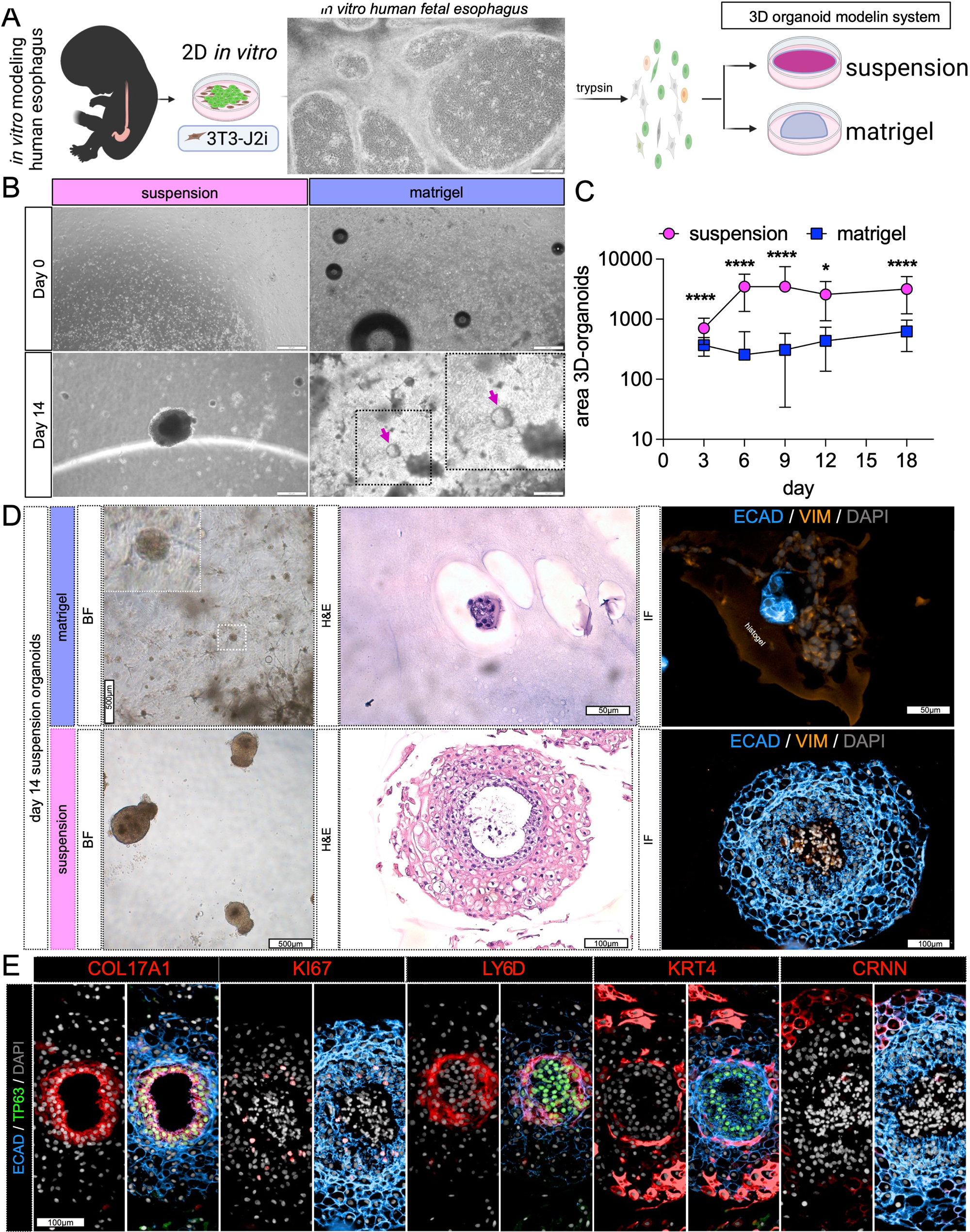
Development of 3D Suspension Organoids from Early Development Esophageal Cells. (**A**) Schematic representation of the in vitro modeling process for the human early development esophagus. Cells isolated from early development esophageal tissues were expanded in 2D on a 3T3-J2 feeder layer and transitioned into 3D culture systems, either in suspension or Matrigel. (**B**) Brightfield images of organoids in suspension and Matrigel systems at Day 0 and Day 14. Suspension organoids exhibit defined spherical morphology, while Matrigel organoids show irregular structures with internal cavities (magenta arrows). Scale bars: 500 µm (Day 0) and 200 µm (Day 14). (**C**) Quantification of the area of 3D organoids over time (Days 3, 6, 9, 12, 15, and 18) shows significant differences in growth between suspension and Matrigel systems. Suspension organoids exhibit a larger size compared to Matrigel organoids. Data are presented as mean ± SEM, *p < 0.05, ****p < 0.0001. (**D**) Brightfield (BF), hematoxylin and eosin (H&E), and immunofluorescence (IF) images of Matrigel and suspension organoids at Day 14. H&E staining shows stratified epithelial layers in suspension. IF staining for ECAD (epithelial marker) and VIM (mesenchymal marker) reveals epithelial organization and mesenchymal components in suspension organoids. Scale bars: 500 µm (BF), 100 µm (H&E), and 50 µm (IF). (**E**) Immunofluorescence staining of suspension organoids at Day 14 for epithelial and basal markers, including COL17A1(basal layer marker), KI67 (proliferation marker), LY6D (early basal marker), KRT4 (suprabasal marker), and CRNN (luminal marker). Organoids display distinct epithelial stratification with spatially localized marker expression. Scale bar: 100 µm.

**Supplemental Figure 3:**
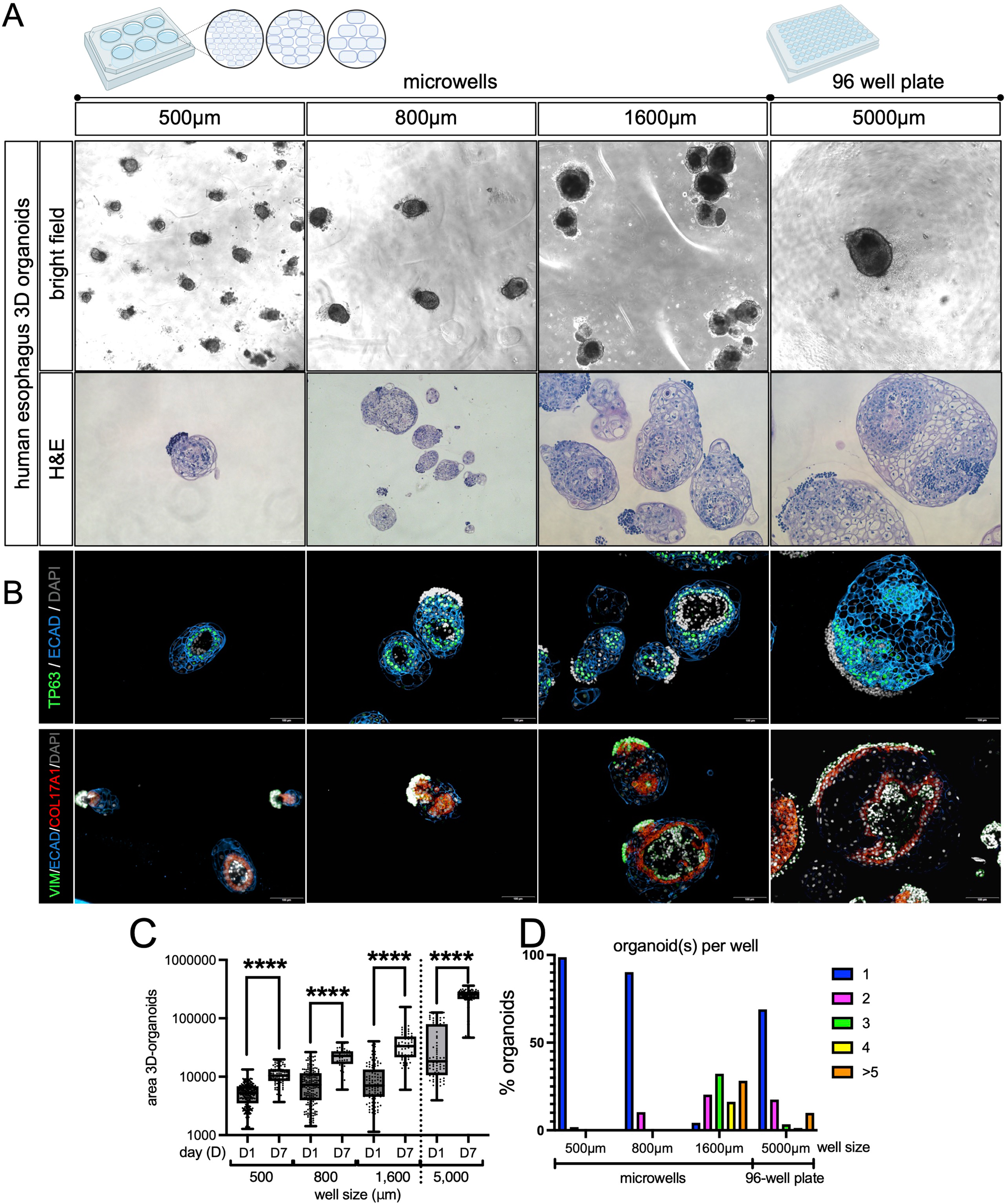
Optimization of Organoid Size and Uniformity Using Microwells. (**A**) Brightfield and hematoxylin and eosin (H&E) images of human esophageal 3D organoids cultured in microwells of varying diameters (500 µm, 800 µm, 1600 µm, and 5000 µm). Smaller microwells (500 µm and 800 µm) produce uniform, compact organoids, while larger microwells (1600 µm and 5000 µm) generate heterogeneous organoids with variable sizes and morphologies. Scale bar: 200 µm (brightfield) and 50 µm (H&E). (**B**) Immunofluorescence staining of organoids cultured in microwells of different sizes. TP63 (green, basal progenitor marker), ECAD (blue, epithelial marker), and COL17A1 (red, basal layer marker) reveal stratified epithelial organization. Larger microwells show increased variability in epithelial layer integrity and cell distribution. Scale bar: 50 µm. (**C**) Quantification of organoid area on Day 1 and Day 7 across different microwell sizes. Smaller microwells (500 µm and 800 µm) produce significantly smaller, more uniform organoids, while larger microwells (1600 µm and 5000 µm) result in greater organoid variability. Data are presented as mean ± SEM, ****p < 0.0001. (**D**) Distribution of organoid numbers per well across different microwell diameters. Smaller microwells (500 µm and 800 µm) predominantly yield single organoids per well, whereas larger microwells frequently generate multiple organoids per well, highlighting the impact of microwell size on organoid uniformity.

